# Most commonly mutated genes in High Grade Serous Ovarian Carcinoma are nonessential for ovarian surface epithelial stem cell transformation

**DOI:** 10.1101/2020.01.27.921718

**Authors:** Robert J. Yamulla, Shreya Nalubola, Andrea Flesken-Nikitin, Alexander Yu. Nikitin, John C. Schimenti

## Abstract

High grade serous ovarian carcinoma (HGSOC) is the most lethal gynecological cancer and the 5^th^ leading cause of cancer-related deaths of women in the USA. Disease-associated mutations have been identified by the Cancer Genome Atlas Research Network. However, aside from mutations in *TP53* or alterations in the *RB1* pathway that are extremely common in HGSOC, the contributions of other mutation combinations have been difficult to assess experimentally or with genomic data alone. Previous research identified ALDH^+^ stem cells of the ovarian surface epithelium (OSE) as one of the putative cells of HGSOC origin. Here, we performed combinatorial CRISPR mutagenesis of 20 putative HGSOC driver genes to identify mutation combinations that transformed OSE stem cells (OSE-SC) and non-stem cells (OSE-NS). Overrepresented mutations and mutation combinations were identified in all transformants and were investigated directly in targeted assays. Our results support the OSE stem cell theory of HGSOC initiation and suggest that most commonly mutated genes in HGSOC have no effect on OSE-SC transformation initiation. We suggest a model in which combined disruption of *RB1* and *PTEN*, in addition to *TP53* deficiency, constitutes a core set of mutations required for efficient transformation *in vitro.* A few previously uncharacterized mutation combinations further enhanced transformation but may have done so via TP53-related mechanisms. Together, our results identify mutation combinations that are critical for OSE-SC transformation and may contribute to more accurate modeling of HGSOC development. Our cancer driver screening methodology may also serve as a model for high throughput functional assessment of commonly mutated genes uncovered in other cancers by large scale sequencing.

## Introduction

Ovarian cancer is a complex disease consisting of several distinct subtypes that differ in progression, prognosis, cell of origin, and genetic alterations (Bowtell 2010). It is the fifth leading cause of cancer-related female deaths in the western world (Siegel et al. 2020). High grade serous ovarian carcinoma (HGSOC) is the most common (about 70%) and lethal subtype of ovarian cancer, in part due to its propensity to metastasize, and to relapse following chemotherapy (Auersperg 2013). Additionally, HGSOC screening methodologies are inefficient, typically resulting in late stage diagnosis. The rarity of early stage HGSOC detection has complicated ascertainment of the cell of origin, initiating mutations, and the identification of precursor lesions (Feeley and Wells 2001; Auersperg 2013).

Much progress has been made regarding the genetic etiologies of HGSOC. The Cancer Genome Atlas (TCGA) completed a comprehensive genomic analysis of patient tumor samples and commonly dysregulated genes and pathways (Table 1) (Cancer Genome Atlas Research Network 2011). Common mutations and deletions of genes are of particular interest, as they may drive HGSOC initiation and development. Several of the putative TCGA driver genes have been thoroughly investigated. For instance, *TP53 (Trp53* in mice) is mutated or inactivated in nearly all tumors (Cancer Genome Atlas Research Network 2011), and has been validated as a crucial driver of carcinogenesis in mouse models (Harlan and Nikitin 2015; Bobbs et al. 2015; Kim et al. 2018).

**Table 1:** Alteration frequency of minilibrary target genes in HGSOC. **Legend:** Putative HGSOC driver genes were derived primarily from the Cancer Genome Atlas Research Network. Most genes, except for *FANCM* and *APC,* were found to be significantly mutated or deleted in HGSOC tumors by TCGA.

| **Gene Name** | **Mutations** | **Amplifications** | **Deletions** | **Total** | **% Disrupted** |
| --- | --- | --- | --- | --- | --- |
| *Trp53* | 303 | 2 | 1 | 306 | 96.20% |
| *Brca1* | 37 | 1 | 1 | 39 | 12.03% |
| *Brca2* | 34 | 3 | 2 | 39 | 11.39% |
| *Csmd3* | 19 | 47 | 1 | 67 | 6.33% |
| *Nf1* | 12 | 1 | 24 | 37 | 11.39% |
| *Fat3* | 18 | 8 | 1 | 27 | 6.01% |
| *Gabra6* | 6 | 3 | 1 | 10 | 2.22% |
| *Rb1* | 6 | 1 | 25 | 32 | 9.81% |
| *Apc* | 7 | 2 | 3 | 12 | 3.16% |
| *Lrp1b* | 13 | 6 | 13 | 32 | 8.23% |
| *Prim2* | 0 | 6 | 2 | 8 | 0.63% |
| *Cdkn2a* | 0 | 1 | 7 | 8 | 2.22% |
| *Crebbp* | 7 | 3 | 10 | 20 | 5.38% |
| *Wwox* | 0 | 2 | 14 | 16 | 4.43% |
| *Ankrd11* | 4 | 1 | 10 | 15 | 4.43% |
| *Map2k4* | 1 | 0 | 11 | 12 | 3.80% |
| *Fancm* | 2 | 3 | 0 | 5 | 0.63% |
| *Fancd2* | 1 | 3 | 0 | 4 | 0.32% |
| *Rad51c* | 0 | 2 | 0 | 2 | 0.00% |
| *Pten* | 2 | 2 | 21 | 25 | 7.28% |

However, most of the recurrently altered HGSOC driver genes are mutated or deleted in a smaller subset of tumors and have not been validated experimentally in animal models or cell transformation paradigms. Several, including *WWOX*, *LRP1B*, *CDKN2A*, and *PTEN*, exist near fragile sites in the genome, and therefore may be mutated simply as a consequence of genome instability rather than cancer initiation (Hess et al. 2019).

Although *TP53* is mutated in nearly all HGSOC cases, experiments with mouse models indicate that *Trp53* mutagenesis is insufficient for HGSOC initiation; rather, multiple mutations appear to be required, consistent with the multi-hit hypothesis of cancer (Knudson 1971; Flesken-Nikitin et al. 2003). For example concurrent inactivation of *Trp53* and *Brca1*, both commonly mutated in HGSOC, could not drive transformation in mice (Xing and Orsulic 2006). However, activation of *Myc* along with disruption of both *Trp53* and *Brca1* did initiate HGSOC. It was also reported that disruption of *Trp53* or *Rb1* alone in the ovarian surface epithelium (OSE) caused neoplasms in only 4/31 and 1/21 mice, respectively, but simultaneous mutation of both caused 100% cancer incidence after a median 227 days (Flesken-Nikitin et al. 2003). Given the substantial numbers of commonly mutated genes identified by TCGA, there are a myriad of possible TCGA driver gene combinations, but very little data regarding how these different combinations could affect transformation efficiency of putative HGSOC “cells of origin.”

There is a growing consensus that HGSOC may have several places of origin, such as OSE, tubal epithelium and peritoneal serosa (Harlan and Nikitin 2015; Kim et al. 2018; Zhang et al. 2019; Lawrenson et al. 2019; Hao et al. 2017). The OSE is a flat to cuboidal cell monolayer that overlies the ovary and was originally proposed as the HGSOC putative cell type-of-origin due to correlation with tumor localization and the observation that a greater number of ovulatory cycles correlates with increased cancer incidence (Fathalla 1971; Fathalla 2013; Okamura et al. 2006; Auersperg 2013).

Research has suggested that repeated cycles of follicular rupture, OSE damage, inflammation, and repair may trigger oncogenic transformation of OSE (Katabuchi and Okamura 2003). Inclusion cysts, or entrapment of OSE within the ovarian stroma, may also facilitate OSE transformation by exposing it to high concentrations of hormones, growth factors, and inflammatory cytokines that are not present at the ovarian surface. OSE within inclusion cysts has been previously shown to express HGSOC markers like PAX8 (Auersperg 2013). Importantly, the OSE has been experimentally shown to transform into HGSOC-like neoplasms (Flesken-Nikitin et al. 2003; Kim et al. 2018; Zhang et al. 2019).

In mice, HGSOC can initiate from OSE stem cells (OSE-SC) (Flesken-Nikitin et al. 2013). Such cells have high levels of Aldehyde dehydrogenase (ALDH) activity and transform more readily than OSE cells with low ALDH activity (ALDH-, non-stem). OSE-SC were found to have increased colony-forming potential in primary culture, greater sphere formation capacity *in vitro*, and an increased ability to proliferate in culture before undergoing senescence (Flesken-Nikitin et al. 2013). The cells also express multipotency markers and readily transform following combined knockout of *Trp53* and *Rb1* in mice. Thus, the ALDH+ OSE stem cell subpopulation is a candidate originating source of HGSOC.

Here, we sought to identify and functionally validate combinations of putative driver genes in HGSOC, and the transformation susceptibility of different ovarian epithelial cell types to combinations of mutations. The 20 candidate genes tested were primarily those that are most commonly mutated in HGSOC (Table 1). Random sets of mutations were induced by infection of mouse OSE stem cells (OSE-SC) and non-stem cells (OSE-NS) with a minilibrary of lentiviruses encoding Cas9 and CRISPR guide RNAs directed against the candidate driver genes. We found that OSE-SC transform more efficiently than OSE-NS, and that only a fraction of commonly mutated HGSOC genes contribute to transformation *in vitro*. In addition to *Trp53* and *Rb1*, mutation of *Pten* was found to be centrally important for transformation of mouse OSE *in vitro*. We also report novel transformation-enhancing mutation combinations and propose a model of core OSE-SC transformation requirements.

## RESULTS

### Strategy for screening candidate HGSOC suppressors and construction of a validated CRISPR-based lentiviral mini-library

We adapted a strategy (Figure 1), analogous to that described by Zender and colleagues (Zender et al. 2006; Zender et al. 2010), to validate candidate tumor suppressors in a sensitized cell type. Sensitized cells do not undergo transformation in culture or in vivo upon transfer to a mouse host but can do so when an additional gene or combination of genes are disrupted. Since *TP53* is mutated in nearly all HGSOCs but *Trp53* mutagenesis alone does not cause spontaneous ovarian tumors in mice (Flesken-Nikitin et al. 2003; Donehower 1996; Donehower 2014), we decided to use *Trp53*+/- cells in an inbred mouse strain background as the sensitized platform. Accordingly, we created and validated a new null *Trp53* allele in strain FVB/NJ as a source of ovarian surface epithelial cells (Figure S1A-D) for all screening experiments. Strain FVB/NJ was chosen because it is neither susceptible nor resistant to spontaneous ovarian lesions (Huang et al. 2008; Mahler et al. 1996).

**Figure 1.**
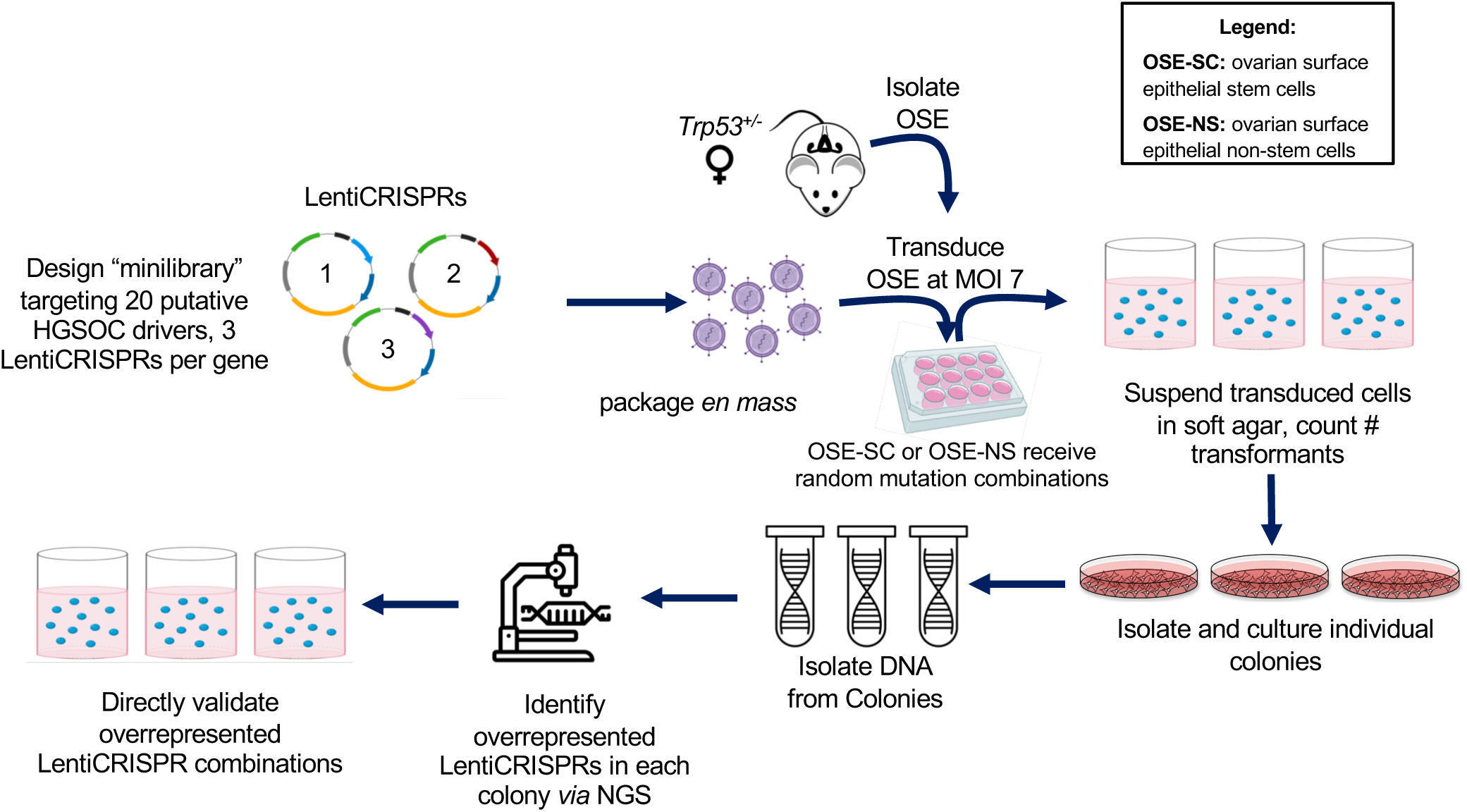
Strategy for identifying HGSOC tumor suppressor combinations. A total of 60 constructs were made in the vector LentiCRISPRv2, constituting the “minilibrary”. OSE-NS or OSE-SC were transduced with functionally-validated LentiCRISPRs and then plated in soft agar. Individual transformants/colonies were isolated and individually cultured. Genome-integrated LentiCRISPRs from each transformant were identified by sequencing, and overrepresented combinations later validated in directed soft agar transformation assays. OSE-SC, ovarian surface epithelium stem cells; OSE-NS, ovarian surface epithelium non-stem cells; NGS, next-generation sequencing.

We generated a list of 20 potential HGSOC driver genes, largely corresponding to the most commonly mutated or deleted genes according to TCGA. We also included *FANCM* due to the putative role of Fanconi anemia-related genes in HGSOC development and *APC* because of to its critical role in canonical Wnt signaling and association with Type I ovarian carcinoma (Cancer Genome Atlas Research Network 2011; Yamulla et al. 2014; Auersperg 2013; Bowtell 2010) (Table 1). We hypothesized that if these genes were true tumor suppressors, then mutating them alone or in combination with *Trp53* and/or other gene mutations would drive transformation in the relevant cell type. We next constructed a lentiviral CRISPR (version 2; hereafter called “LentiCRISPR”) (Shalem et al. 2014; Sanjana et al. 2014) “minilibrary” targeting the 20 putative HGSOC tumor drivers, infected the potential cells-of-origin at a high multiplicity of infection (MOI), and assessed which vectors were overrepresented in transformed cells (See Methods, Figure 1). The library contained 3 constructs per gene, with the sgRNAs in each vector targeting the earliest possible exon of each gene to increase the likelihood of causing loss-of-function indel mutations via error-prone non-homologous end-joining (NHEJ) repair. We tested the minilibrary for gene editing efficiency in pilot assays prior to its use in HGSOC driver screening (Figure S2, Table S1), and designed new guides to replace vectors that did not efficiently mutate targets (Figure S3, Table S2).

### Random combinatorial mutagenesis of candidate HGSOC driver genes in OSE-SC and OSE-NS

To assess transformation frequency *in vitro*, we infected FACS-sorted *Trp53+/-* OSE-SC and OSE-NS (Figure S4) with either the lentiCRISPR minilibrary or a GFP-containing lentivirus, both at an MOI of ∼ 7 (Figures S5, S6). The cells were then plated in soft agar to allow for assessment of adhesion-independent growth, a hallmark of carcinogenesis and transformation (hereafter, formation of adhesion independent colonies will be referred to as “transformation”, and individual colonies will be referred to as “transformants”) (Horibata et al. 2015; Borowicz et al. 2014; Puck et al. 1956; Roberts et al. 1985; de Larco and Todaro 1978; Hamburger and Salmon 1977). No colonies were observed in either non-transduced cells or cells infected with the GFP-containing vector (Figures S1E,F; S6; S7). Thus, heterozygosity for *Trp53* alone, or the process of infection, did not enable transformation of either cell type. However, transformants were observed in cells transfected with the entire library, albeit at low frequency despite an efficient infection frequency as judged by control infections with GFP lentivirus and LentiCRISPRv2 serial dilution assays (Figures 2A; S5, S7). OSE-SC were transformed (formed colonies) 41-fold more frequently than OSE-NS or unsorted OSE (0.66%, 0.02% and 0.21% of total plated cells, respectively) (Figure 2A, S7). The low transformation frequency suggests that only a few mutation combinations can initiate colony growth. That unsorted OSE produced 10.5-fold more colonies than OSE-NS is likely due to its OSE-SC subpopulation (Figures 2A; S4A, S7). To rule out the possibility that OSE-SC are simply infected by lentiviruses more efficiently than OSE-NS, we transduced each cell type with equal concentrations of mCherry lentivirus and scored for mCherry expression in each population. OSE-SC and OSE-NS were transduced at similar rates (72.9% and 85.6% mCherry expression, respectively), suggesting that transformation efficiency differences were indeed cell-type specific (Figure 2B,C).

**Figure 2.**
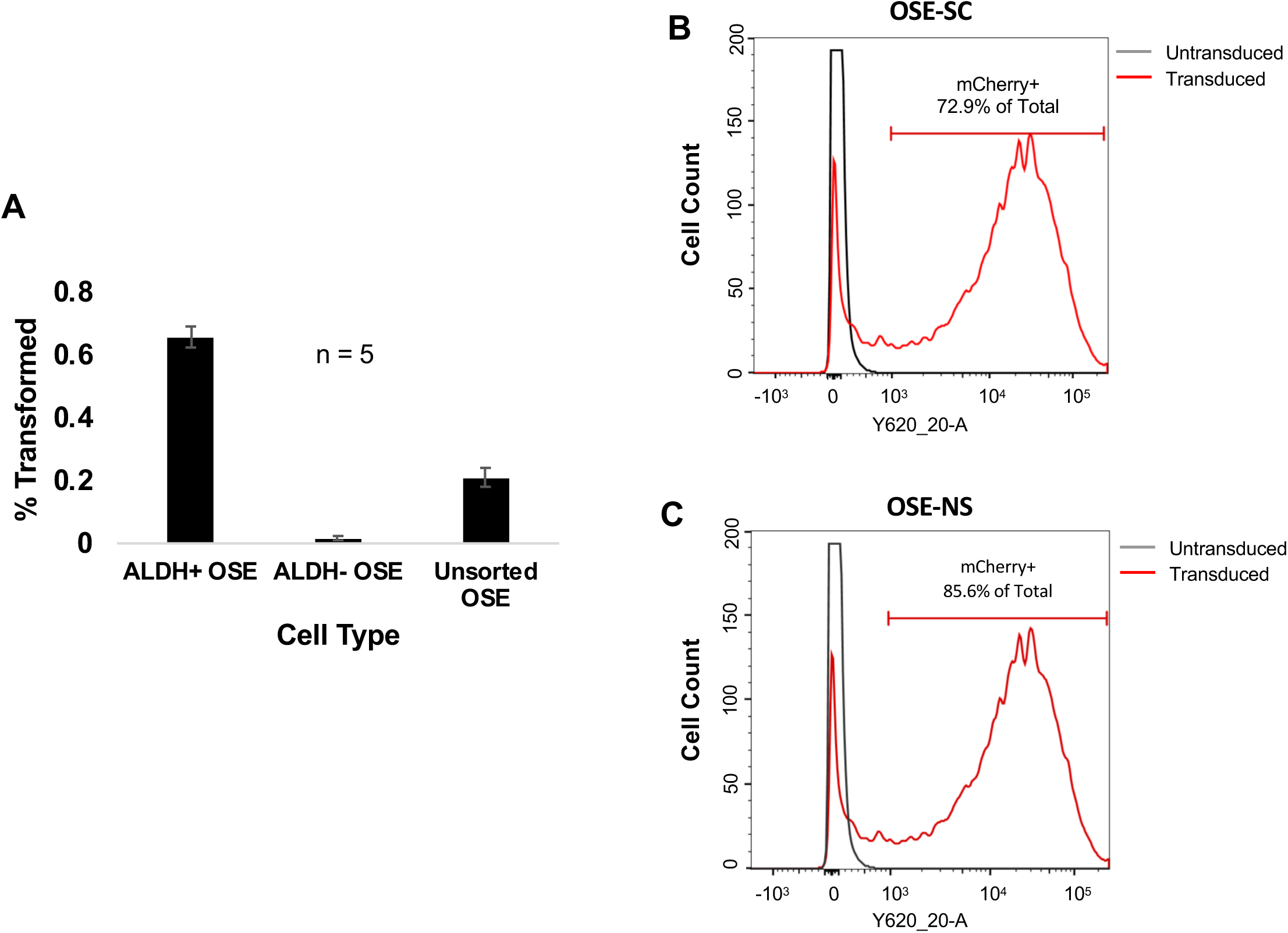
OSE-SC (ALDH+) transform more frequently than OSE-NS (ALDH-) despite similar viral transduction rates. **(A)** Percent transformation of OSE-SC, OSE-NS and unsorted OSE following LentiCRISPRv2 minilibrary transduction. OSE-SC transformed more frequently than OSE-NS and unsorted OSE. Unsorted OSE transformed more frequently than OSE-NS (5 replicates. SEM error bars). (**B,C)** FUGW-mCherry (mCherry-expressing lentivirus) transduction efficiency in OSE-SC and OSE-NS detected via flow cytometry. Flow cytometry was used to count mCherry+ cells following transduction with equal concentrations of FUGW-mCherry lentivirus. Percentages indicate the percentage of total cells that are mCherry+. The dark grey lines represent cell counts of untransduced cells. The red line represents cell counts of FUGW-mCherry transduced cells. OSE-SC and OSE-NS gained mCherry fluorescence at similar rates following lentiviral transduction.

Given an MOI of 7, and 3 LentiCRISPRs per gene, each gene represented in the minilibrary should be present in 30.2% of transduced cells (Figure S8). Control experiments supported this estimate and lack of technical biases, including the appearance of a control GFP LentiCRISPR at an expected random frequency of 12% (Methods; Figures 3A,B; S8).

**Figure 3.**
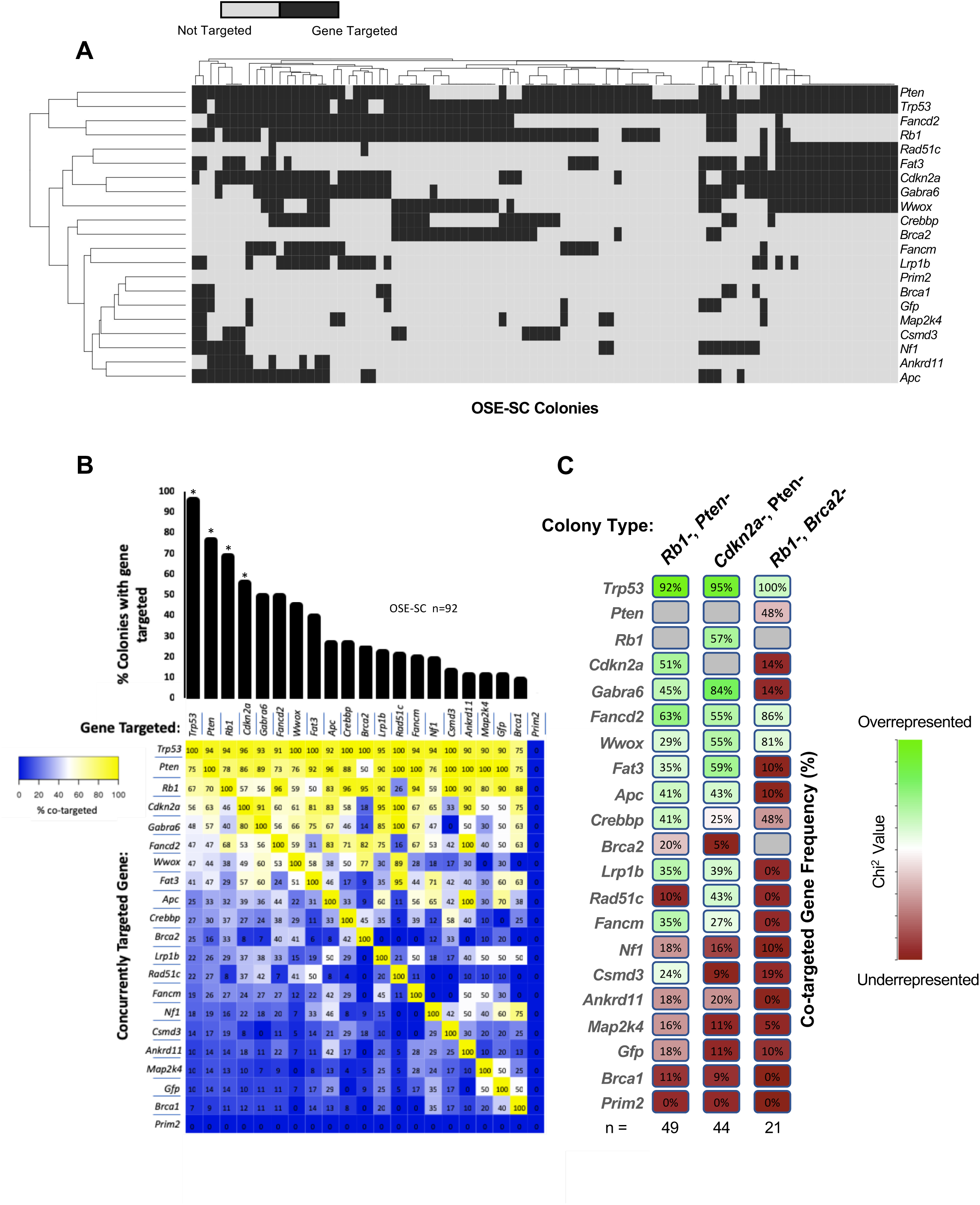
Identification of genome-integrated LentiCRISPRs and overrepresented target gene combinations in OSE-SC. **(A)** Genome-integration and hierarchical clustering of LentiCRISPRv2 constructs in OSE-SC samples. Hierarchical clustering was performed on both sample similarity and gene targeting. **(B)** Overall percent gene targeting and co-targeting frequency. Significance for single integration overall was assessed using Chi2 (df=19) and is indicated with an asterisk. The heat map displays co-integration frequency of each gene present on the x axis with a gene shown on the y axis. **(C)** Overrepresentation of co-targeted genes in sample subgroups. Over or underrepresentation was determined using Chi2 analyses. Chi2 values corresponding to p ≤ 0.05 (df = 19) are colored in green. Red coloration indicates that co-integration may have occurred by chance, and that the p value is ≥ 0.05.

### Enrichment of mutated gene combinations in transformed OSE-SC samples

Next, we determined whether particular minilibrary constructs and combinations thereof were over-represented in the transformants. To do this, we isolated all individual colonies and cultured them as transformed, adherent, clonal cell lines. The LentiCRISPR vectors were identical save for a unique 20bp Cas9 guide sequence which provided a unique molecular barcode (Table S3). We therefore identified all genome-integrated LentiCRISPRs from genomic DNA isolated from each transformant line using a next-generation sequencing-based approach (Figure 1). Three LentiCRISPRs per target gene were included in our minilibrary to control for unintended effects of any single construct. Overrepresented integration of only one of three gene targeting constructs would indicate potential spurious technical errors such as excessively high titer or off-target effects. Our sequencing dataset revealed genome integration of all three LentiCRISPRs corresponding to target genes found in more than 30% of colonies, suggesting that gene targeting was not a consequence of technical issues (Figure S9A).

### *Trp53* loss alone is necessary but insufficient for OSE-SC transformation

Nearly all (96%) OSE-SC colonies contained LentiCRISPRs targeting *Trp53*, consistent with human tumor samples which almost universally harbor *TP53* mutations (Figure 3A,B) (*X*2 (19); p < 0.05). As expected given the high MOI, most OSE-SC transformants (88 of 92) also harbored lentiCRIPSRs targeting other genes. The most common genome-integrated vectors corresponded to *Pten* (76%), *Rb1* (68%), *Cdkn2a* (55%), *Fancd2* (49%), *Wwox* (45%), *Gabra6* (49%), *Fat3* (39%), *Apc* (26%), *Crebbp* (26%), and *Brca2* (24%) (Figures 3A,B; S9B). The rarity of transformants lacking *Trp53* lentiCRISPRs (4%) suggests that heterozygosity of *Trp53* is insufficient for transformation, even in the context of other mutations that can synergize with partial *Trp53* loss (Figure 3A,B). Frequent co-targeting of *Trp53* also supports previous assertions that inactivating mutations in *Trp53* are crucial transformation precursors but are alone insufficient for OSE transformation (Flesken-Nikitin et al. 2003).

### LentiCRISPRs targeting *Trp53*, *Rb1*, *Cdkn2a*, and *Pten* are overrepresented in OSE-SC transformants

In addition to *Trp53,* most colonies (74%) had LentiCRISPRs targeting either *Rb1* or *Pten*, and nearly half (45%) had LentiCRISPRs targeting both (Figure 3 A,B). Most *Pten-* colonies (hereafter, we will refer to transformants bearing particular LentiCRISPRs by the corresponding gene symbol followed by “-“; e.g. *Pten1-*, with the caveat that the actual target gene may not have been mutated to a null state) were also *Rb1-* (70%), suggesting that concurrent inactivation of these two genes facilitates transformation (Figure 3B). This observation supports significant co-occurrence of *RB1* and *PTEN* mutations in HGOSC, and also in many other cancers, such as metastatic prostate cancer, lipomas, and astrocytomas (Gao et al. 2013; Cerami et al. 2012; Cancer Genome Atlas Research Network 2011; Hamid et al. 2019; Filtz et al. 2015; Chow et al. 2011). Although *Rb1* was not targeted in all *Pten-* colonies, 88% of *Pten-* colonies lacking a *Rb1* LentiCRISPR had at least one LentiCRISPR targeting *Cdkn2a* (Figure 3A), which encodes p14ARF and p16INK4a. These are important regulators of TRP53 and RB1, respectively (Zhao et al. 2016; Li et al. 2011; Nielsen et al. 1998). Only 3% of *Pten-* colonies in our dataset had no *Rb1* or *Cdkn2a* LentiCRISPRs, suggesting that either direct or indirect disruption of *Rb1* is important in *Pten-* colonies (Figure 3A).

### Mutations in *Trp53*, *Rb1*, *Pten*, and *Cdkn2a* function synergistically to promote transformation

To validate and dissect the apparent major contributions of *Trp53*, *Pten*, *Rb1* and *Cdkn2a* in suppressing OSE-SC transformation, we performed additional infections with vectors corresponding to combinations of just these 4 genes. Confirming our observations from the whole library screen (see above), mutating the second allele of *Trp53* alone in *Trp53+/-* OSE-SC was inefficient in transformation (Figures 4A; S10, S11A). However, co-mutagenesis with combinations of *Rb1*, *Cdkn2a*, and *Pten* significantly enhanced transformation efficiency (colony number) and colony size (Figures 4A; S10; S11A). Specifically, we observed that targeting *Trp53* alongside *Pten* and *Rb1*, *Pten* and *Cdkn2a*, or *Pten*, *Rb* and *Cdkn2a* led to greater transformation efficiencies and larger colonies (Figures 4A; S10; S11A). The most efficient transformation and largest colony sizes occurred when *Trp53, Rb1, Cdkn2a*, and *Pten* were targeted simultaneously (Figure 4A; S10; S11A). There was no significant increase in transformation efficiency when *Trp53* was co-targeted with only one of the other 3 genes, although there was a significant increase in the size of *Trp53-/Rb1-* and *Trp53-/Cdkn2a-* colonies (Figure S11A). These results indicate that deficiency of *Trp53, Rb1* and *Pten* constitute a core state for efficient transformation of OSE-SC *in vitro*.

**Figure 4.**
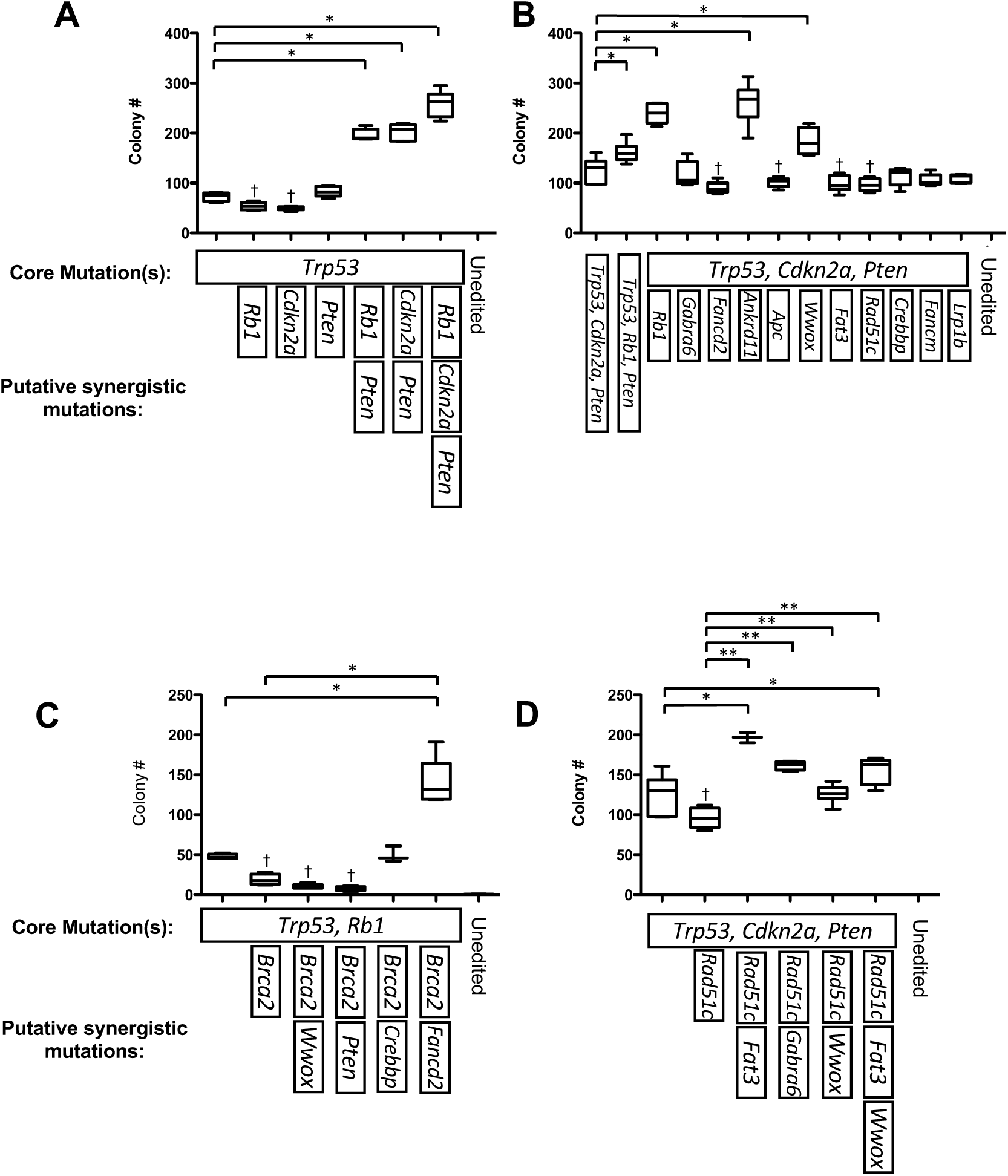
Targeted OSE-SC-transformation assay and validation of overrepresented LentiCRISPR combinations. A baseline level of adhesion independent growth was first assessed via induction of specific “core mutations” via LentiCRISPRv2 targeting. Additional minilibrary target genes were then mutated (using LentiCRISPRv2) alongside core mutations to assess whether they act synergistically to promote adhesion independent growth. Colony counts that are significantly greater than baseline rates (core mutations alone) are labeled with an asterisk (*) (Students’ two-tailed t-test p<0.05). Colony counts that are significantly lower than baseline rates are labeled with an obelisk (†). Standard error of the mean (SEM) error bars. **(A)** Targeted transduction of *Trp53*, *Rb1*, *Pten* and *Cdkn2a* LentiCRISPRs in OSE-SC. Significantly greater rates of OSE-SC transformation vs OSE-SC transduced with *Trp53* LentiCRISPRs alone occurred when all four genes were targeted together, or via mutagenesis of *Trp53*, *Rb1* (or *Cdkn2a*) and *Pten* (Students’ two-tailed t-test p ≤ 0.05, n=6). **(B)** Targeted transduction of *Trp53*, *Cdkn2a* and *Pten* LentiCRISPRs plus putative transformation enhancers. Only the addition of *Ankrd11* or *Wwox* LentiCRISPRs to *Trp53*, *Cdkn2a* and *Pten* LentiCRISPRs significantly enhanced colony formation vs *Trp53-/Cdkn2a-/Pten-* OSE-SC (Students’ two-tailed t-test p<0.05, n=6). The addition of *Fancd2*, *Apc*, *Fat3*, and *Rad51c* significantly decreased colony count (Students’ two-tailed t-test p<0.05). **(C)** Targeted transduction of *Brca2* LentiCRISPRs and *Brca2*-associated LentiCRISPRs. *Fancd2* mutations functioned synergistically with *Brca2, Trp53 and Rb1 mutations* to significantly enhance colony formation vs *Trp53-*/*Rb1-*/*Brca2-* or *Trp53-*/*Rb1-* OSE-SC (Students’ two-tailed t-test p ≤ 0.05, n=6). **(D)** Targeted mutagenesis of *Rad51c* LentiCRISPRs and *Rad51c-*associated LentiCRISPRs. *Fat3* and *Gabra6* mutagenesis alongside *Trp53*, *Cdkn2a*, *Pten* and *Rad51c* significantly increased colony count compared to *Trp53-/Rb1-/Pten-/Rad51c-* OSE-SC (Students’ two-tailed t-test p ≤ 0.05, n=6).

### Disruption of *Ankrd11* or *Wwox* can further enhance transformation of *Trp53*-/*Cdkn2a*-/*Pten*-OSE-SC

Interestingly, certain cohorts of genes were co-targeted at significant frequencies in *Trp53*-/*Rb1*-/*Pten-* (*Cdkn2a*, *Gabra6*, *Fancd2, Wwox*, *Fat3*, *Apc*, *Crebbp*, *Lrp1b*, *Fancm*, and *Csmd3*; *X*2 = 19, p < 0.05) or *Trp53*-/*Cdkn2a-*/*Pten-* colonies (*Rb1*, *Gabra6*, *Fancd2*, *Wwox*, *Fat3*, *Apc*, *Lrp1b*, *Rad51c*, and *Fancm*; *X*2 = 19, p < 0.05), despite vectors for each being underrepresented overall (Figure 3B,C). To explore whether mutations in these genes influenced transformation, we performed additional OSE-SC infections in which individual genes were mutated along with the core combinations of *Trp53*, *Cdkn2a* (or *Rb1*) and *Pten*. Most (*Lrp1b*, *Fancm*, *Crebbp*, *Rad51c*, *Fat3*, *Apc*, *Fancd2*, and *Gabra6*) did not enhance transformation rates, and several (*Fancd2*, *Apc*, *Fat3*, and *Rad51c*) actually decreased transformation rates (Figures 4B; S12).

However, significant increases in transformation frequency were observed when LentiCRISPRs targeting *Ankrd11* or *Wwox* were added to the core of *Trp53, Cdkn2a* and *Pten* (Figure 4B). No significant increase in colony size was noted following mutagenesis of any TCGA driver genes other than *Trp53*, *Rb1*, *Cdkn2a*, and *Pten* (Figure S11B). We surmised that the overall underrepresented target genes that had no (or a negative) effect on OSE-SC transformation in targeted experiments, but which were often present in cells with high numbers of other vectors, was a technical artifact. Indeed, a correlative relationship between underrepresented target genes and the number of genes targeted per colony (Figure S13). For instance, most samples with LentiCRISPRs targeting *Ankrd11*, *Apc*, *Lrp1b*, *Brca1*, *Nf1*, *Fancm, Fancd2*, and *Map2k4* occurred in colonies with 8 or more targeted genes (Figure S13). Such clones also generally contained common LentiCRISPRs for *Trp53*, *Rb1*, *Pten*, and *Cdkn2a* (Figure 3A,B), suggesting that many or all of these lower-frequency “hits” are unrelated to transformation, but perhaps the parental cell was particularly susceptible to viral infection.

### *Brca2* disruption deters transformation of *Trp53*-/*Rb1*-OSE-SC, but enhances transformation when co-mutated with *Trp53*, *Rb1*, and *Fancd2*

Notably, LentiCRISPRs for *Brca2* were underrepresented overall in OSE-SC colonies, despite the association of mutations in this gene with familial HGSOC (Figure 3 A,B) (Risch et al. 2006; Walsh 2015; Cancer Genome Atlas Research Network 2011). *Brca2* deficiency is cell lethal in the absence of other mutations, causing replication stress, mitotic abnormalities, 53BP1 activation, and G1 arrest (Feng and Jasin 2017; Zhu et al. 2015). However, most cancer cells develop mechanisms to overcome this G1 arrest through rescuing mutations in other genes like *Trp53* (Feng and Jasin 2017). In our screen, 95% of *Brca2*-colonies were also *Trp53*-/*Rb1*- (Figure 3A,B). Many *Brca2*-colonies also contained LentiCRISPRs targeting *Fancd2* (82%), *Wwox* (77%), and *Crebbp* (45%) (Figure 3B). Constructs targeting *Trp53*, *Fancd2* and *Wwox*, in particular, were overrepresented alongside *Brca2* and *Rb1* (*X*2 (19); p < 0.05). We hypothesized, therefore, that many of these co-targeted genes are necessary for efficient *Brca2*-colony growth. Targeted LentiCRISPR co-infections revealed that *Brca2* mutation significantly reduced transformation rates of cells also targeted for *Trp53* and *Rb1*, in agreement with previous reports (Figure 4C; S14) (Feng and Jasin 2017; Zhu et al. 2015). However, we found that the addition of *Fancd2* LentiCRISPRs to constructs targeting *Trp53*, *Rb1* and *Brca2* rescued the detrimental effects of single mutations in either *Brca2* or *Fancd2* on transformation rate, and significantly increased colony size (Figure 4C; S11C, S14). *Trp53*-/*Rb1*-/*Fancd2*-/*Brca2*-cells had 3 fold more colonies than *Trp53*-/*Rb1*-OSE-SC (Figure 4C; S14). Despite co-targeting with *Brca2* in 48% of cases, neither *Pten* nor *Wwox* LentiCRISPR transductions significantly increased number or size relative to *Trp53*-/*Rb1*- or *Trp53-*/*Rb1*-/*Brca2*-colonies (Figure 4C; S11C, S14). We did, however, find that *Crebbp* targeting restored *Trp53-*/*Rb1*-/*Brca2*-colony formation to the level observed for *Trp53*-/*Rb1*-colonies, and significantly increased colony size (Figure 4C; S11C). These results suggest that multiple concurrent driver mutations are necessary to overcome growth-deterring effects of *Brca2* mutagenesis.

### *Rad51c* synergizes with *Fat3* and *Gabra6* to promote transformation

Like *Brca2*, *Rad51c* is involved in DNA double strand break repair (Somyajit et al. 2010). Loss of *Rad51c* has also been shown to be detrimental to cell growth, so synergistic mutations may be required for efficient adhesion independent growth of *Rad51c*-OSE-SC (Kuznetsov et al. 2009). Although *Rad51c* mutagenesis was detrimental to the transformation of *Trp53*-/*Cdkn2a*-/*Pten*-OSE-SC (Figure 4B), we observed that *Rad51c* was frequently co-mutated with *Gabra6* (100%), *Wwox* (89%) and *Fat3* (95%) in our screen (Figures 3B). We therefore performed targeted infections of LentiCRISPRs targeting *Gabra6*, *Wwox*, and *Fat3* alongside *Rad51c* and core mutations in *Trp53*, *Cdkn2a*, and *Pten*. We observed significantly increased transformation rates following concurrent mutagenesis of *Rad51c* and *Fat3* or *Gabra6*, but not *Wwox*, in *Trp53*-/*Cdkn2a*-/*Pten*-colonies (Figure 4D; S15). Only *Trp53-/Cdkn2a-/Pten-/Rad51c-/Gabra6-* colonies were significantly larger than *Trp53-/Cdkn2a-/Pten-* or *Trp53-/Cdkn2a-/Pten-/Rad51c-* colonies (Figure S11D). These results suggest that some HGSOC-associated mutations, like those in *Rad51c*, can promote transformation only with additional synergistic mutations to overcome synthetic lethality.

### LentiCRISPR integration patterns in OSE-NS transformants

OSE-NS transformed much less efficiently than OSE-SC, yielding only 11 colonies for analyses (Figure 5). This small sample size preempted meaningful statistical analyses of target gene overrepresentation. Nevertheless, like OSE-SC, most (10/11) clones contained LentiCRISPRs targeting *Trp53*, *Rb1*, *Pten*, and *Cdkn2a*. Other overall commonly targeted genes include *Nf1* (91%), *Crebbp* (82%), *Brca2* (55%), *Brca1* (46%), and *Wwox* (46%).

**Figure 5.**
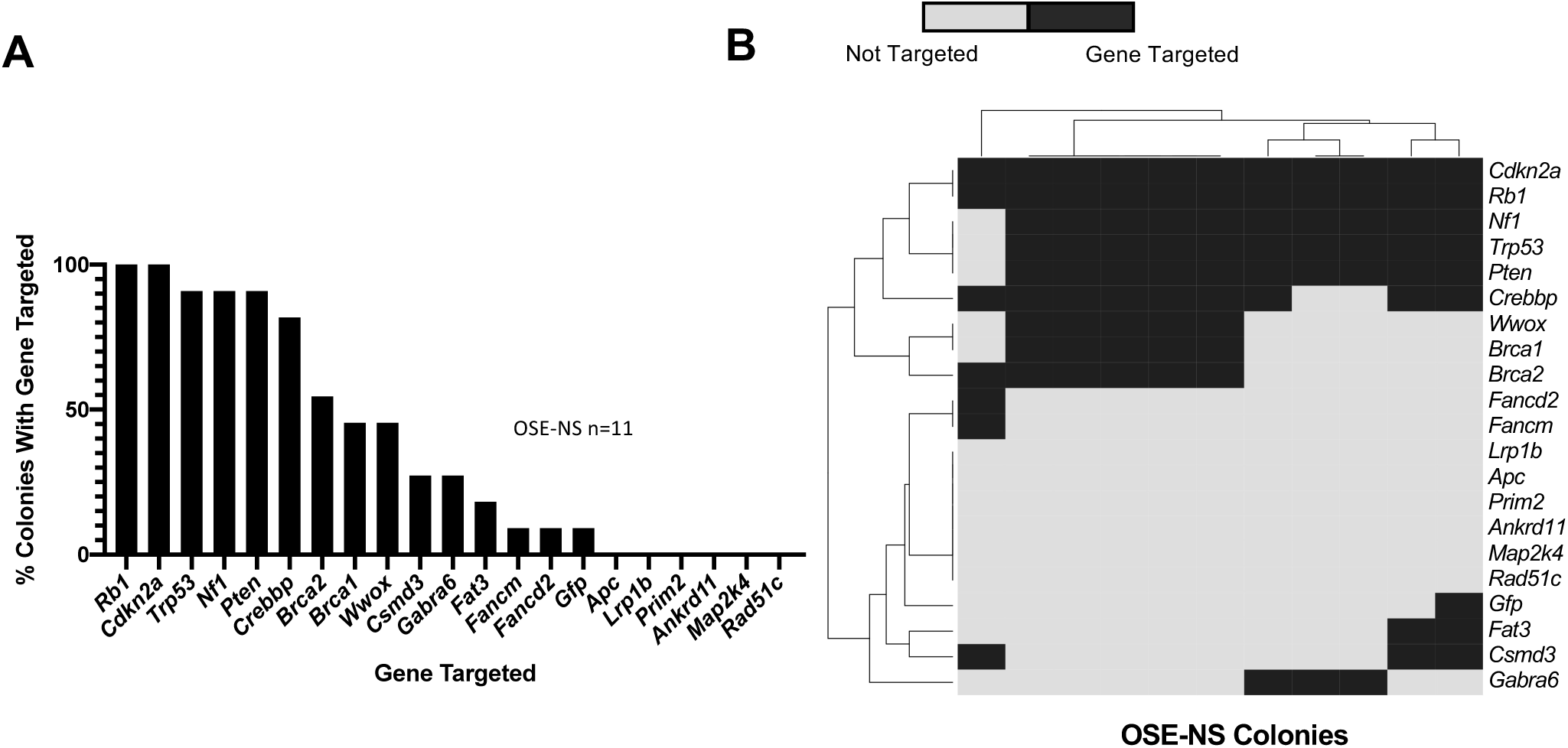
Identification of genome-integrated LentiCRISPRs overrepresented target gene combinations in OSE-NS. **(A)** Percent gene targeting frequency in OSE-NS colonies. **(B)** Genome-integration and hierarchical clustering of LentiCRISPRv2 constructs in OSE-SC samples. The binary color scale shows whether a gene is targeted by at least one lentiCRISPR in each individual sample. Light grey indicates that a given gene was not targeted, while dark grey indicates that a gene was targeted by at least one LentiCRISPRv2 construct. Hierarchical clustering was performed on both sample similarity and gene targeting, resulting in several clusters of co-targeted genes and similar transformants.

## DISCUSSION

It has long been recognized that carcinogenesis typically requires multiple genetic events, and tumor sequencing, as exemplified by the TCGA, has not only supported this tenet, but also informed the constellations of mutations that are commonly present and thus likely contributing to cancer formation and progression. However, the requirements for tumor initiation can be complex in terms of gene combinations and susceptible cells; for HGSOC, precursor lesions have not been conclusively identified, and patient tumor samples have an average of 46 mutations (Cancer Genome Atlas Research Network 2011). Mathematical modeling of cancer driver events suggest that only 5 to 8 driver mutations may be necessary for initiation (Stratton et al. 2009; Pon and Marra 2015), so most of the mutations present in late stage tumors are irrelevant to initiating events.

Whereas contributions of some putative initiating driver genes have been assessed using *in vitro* and *in vivo* systems (Harlan and Nikitin 2015; Hasan et al. 2015; Bobbs et al. 2015; Kim et al. 2018; Zhang et al. 2019), these directed approaches cannot experimentally assess the thousands of mutation combinations in the many commonly altered genes in HGSOC. Our screen represents the first effort to experimentally define the combinations of mutations that drive transformation of ovarian epithelial cells in a random, unbiased manner.

The issue of HGSOC initiation is further complicated by several potential sources of the HGSOC cell of origin. We focused on OSE as a source of putative HGSOC cells of origin, but there are others, such as the distal fallopian tubal epithelium (TE), which can be transformed (Piek et al. 2001; Sherman-Baust et al. 2014; Corzo et al. 2017; Karst et al. 2011; Perets et al. 2013; Labidi-Galy et al. 2017). The presence of serous tubal intraepithelial carcinomas (STICs) in patients with HGSOC, and frequent presence therein of identical *TP53* mutations with those in concurrent HGSOC tumors, led to the proposal that HGSOC can initiate at the TE (Piek et al. 2001; Kindelberger et al. 2007). The TE also shares several well-characterized markers of HGSOC that have not been observed in the OSE (Perets and Drapkin 2016). For example, secretory cells in the TE express PAX8, which is present in HGSOC but not in untransformed OSE (Adler et al.2015; Tacha et al. 2011; Ozcan et al. 2011). Monolayers of PAX8-positive TE cells can also form lesions that express “p53 signatures” that are often associated with mutant *TP53*, but generally have a low proliferative index and lack cellular atypia (Lee et al. 2007; Leonhardt et al. 2011). Correlative links between the TE and HGSOC have been supported by recent algorithmic comparison of HGSOC and distal TE global gene expression landscapes in which similarities were found between HGSOC and the distal TE, but also with OSE-SC in many cases (Lawrenson et al. 2019; Hao et al. 2017). It is possible therefore that all three cell types (OSE-SC, OSE-NS, and TE) have the ability to transform into HGSOC, as has been suggested by others (Neel et al. 2018; Zhang et al. 2019; Kim et al. 2018). These data are supported by direct clinical evidence in which about half of HGSOCs can be reliably explained by STIC origin (Kindelberger et al. 2007; Piek et al. 2001; Auersperg et al. 2008).

We focused on the OSE here because it consists of a single cell type (unlike the heterogeneous TE) for which well-established isolation and culturing methodology exists (Auersperg et al. 2001; Flesken-Nikitin et al. 2003). It is also been shown to transform in animal models such as rat and mouse, and is where HGSOC localizes in patients (Godwin et al. 1992; Testa et al. 1994; Orsulic et al. 2002; Auersperg et al. 2008; Scully 1999; Flesken-Nikitin et al. 2003; Kim et al. 2018). Similarities between OSE-sourced tumors and HGSOC have been demonstrated both genomically and histopathologically (Rosen et al. 2009; Matz et al. 2017; Hao et al. 2017; Zhang et al. 2019; Flesken-Nikitin et al. 2013; Flesken-Nikitin et al. 2003). However, recent evidence has suggested that only a small subpopulation of the OSE (OSE-SC) is particularly cancer-prone (Flesken-Nikitin et al. 2013). Such cells have been shown to possess stem cell-like characteristics such as the ability to replace OSE lost during ovulation and expression of stem cell markers including LGR5, ALDH1, LEF1, CK6B, and CD133 (Flesken-Nikitin et al. 2013). That study also found that *Trp53* and *Rb1* knockouts in OSE-SC result in significantly more tumors and much lower latency compared to *Trp53* and *Rb1* knockouts in non-stem cells.

Our screening results and targeted mutagenesis assays support the theory that the ovarian hilum, which is the putative location of OSE-SC and a transition zone between the OSE, TE, and mesothelium, is particularly prone to transformation. Following random mutagenesis of 20 putative HGSOC driver genes, we found that OSE-SC transformed 41-fold more frequently than OSE-NS. These data support longstanding suspicion that adult stem cells within transition zones are especially prone to carcinogenesis (Ferraro et al. 2010; Greene et al. 1991; Mcnairn and Guasch 2011).

Interestingly, the stem cell theory of cancer initiation may be a unifying concept between OSE- and TE-sourced tumors. Seidman and colleagues (2015) recently demonstrated that most STICs occur in close proximity to the tubal-peritoneal junction (Seidman 2015). Other studies have demonstrated that the tubal-peritoneal junction, like the ovarian hilum region, contains LEF1-expressing cells, and that patients with higher LEF1 expression had poorer 5 year survival (Schmoeckel et al. 2017). It is possible, therefore, that HGSC lesions arise in either OSE-SC or TE stem cells located in transitional zones.

Based on the most common mutation combinations we observed in colonies produced by infection of OSE-SC with the LentiCRISPR minilibrary, and in follow-up validation experiments with specific LentiCRIPSR combinations, we developed a model of events for OSE-SC transformation initiation (Figure 6). We propose that functional loss of only 3 of the 20 genes assessed in this study, *Trp53*, *Rb1*, and *Pten,* are necessary for efficient OSE-SC transformation (Figures 3A,C. 4A; S11A). Knockout of these three genes in the OSE has been previously shown to cause development of both low grade and high grade serous carcinoma in mouse models (Shi et al. 2020).

**Figure 6.**
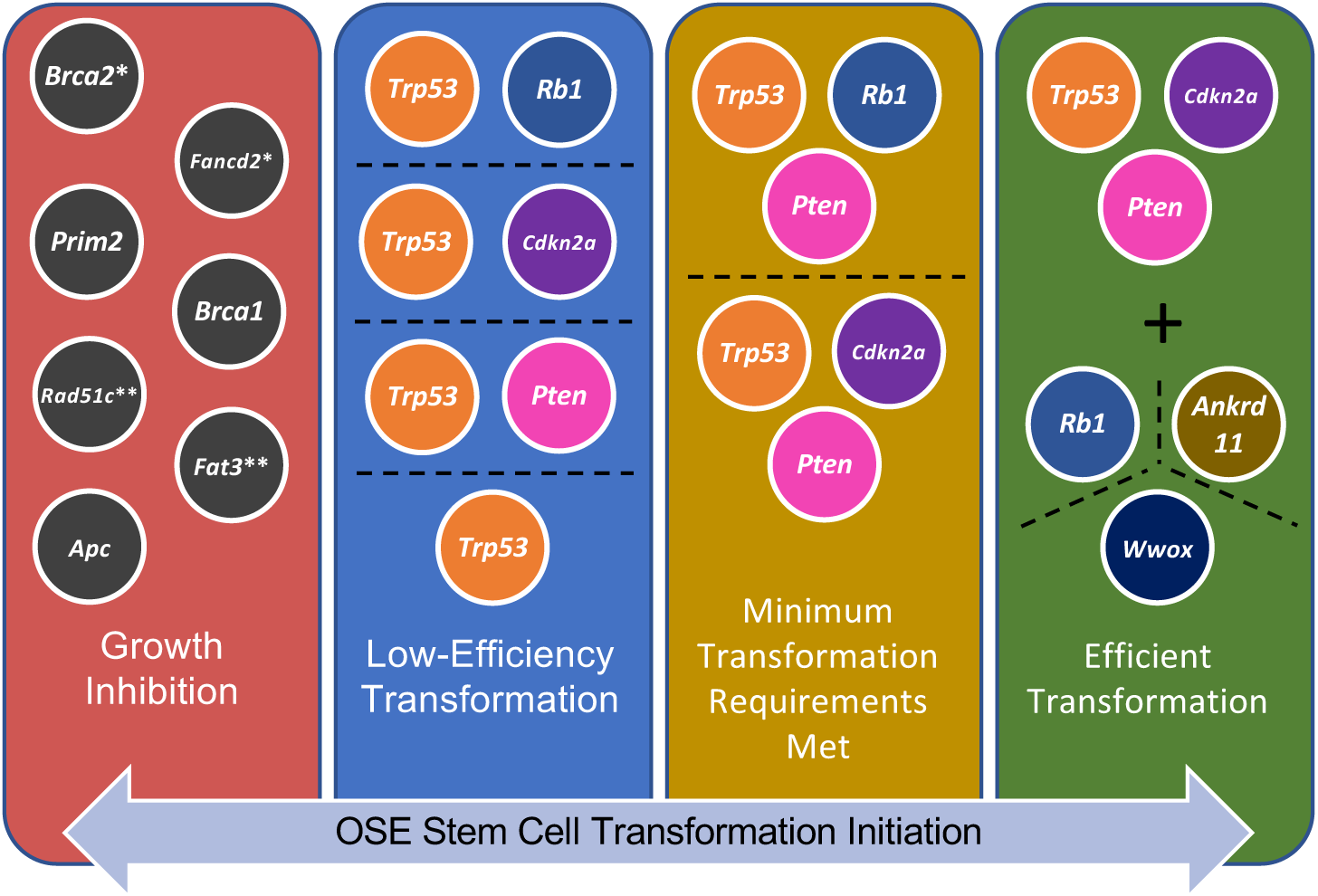
Model of mutations necessary for efficient *in vitro* OSE-SC transformation. Random mutagenesis assays and targeted experiments revealed minimal requirements for adhesion independent growth and mutations that enhance transformation. The blue box contains genes that are ”minimal requirements” for transformation, or cause transformation at low efficiency. The addition of mutations shown in the yellow box cause significant degrees of transformation. Addition of further mutations in genes shown in the green box allow for the highest rates of transformation. Genes listed in the red box inhibit transformation. However, two exceptions exist. *Brca2* and *Fancd2* (marked with a single asterisk) co-mutagenesis alongside *Trp53* and *Rb1* result in efficient OSE-SC transformation. Similarly, *Rad51c* and *Fat3* (or *Gabra6*) (marked with two asterisks) plus *Trp53*, *Cdkn2a* and *Pten* caused efficient transformation.

Mutation of *Cdkn2a* could partially compensate for *Rb1* disruption because *Cdkn2a* encodes p16(INK4a) and p14(ARF), known regulators of *Rb1* and *Trp53*, respectively (Nielsen et al. 1998). Our model also suggests that *Trp53* and/or *Rb1* disruption can cause low efficiency transformation without *Pten* mutations, but colony size was much smaller. The greatest colony size and quantity was observed in *Trp53*-/*Rb1*-/*Cdkn2a*-/*Pten*-colonies, suggesting additive effects of mutations in each gene.

Our proposition that *Trp53* and *Rb1* mutations are core minimal OSE transformation requirements is supported by previous evidence in genetically engineered mouse models (GEMMs). Flesken-Nikitin and colleagues (2003) showed that combined knockout of *Trp53* and *Rb1* in the OSE causes higher rates of carcinogenesis and lower latency compared to single knockout of *Trp53* or *Rb1* alone (Flesken-Nikitin et al. 2003). These results were backed by mouse modeling of *Trp53* and *Rb1* knockouts in the OSE. For example, *Brca1*/*2* or *Trp53* mutations alone failed to cause significant pathologic changes to the OSE, but *Trp53*-/- *Rb1*-/-, *Trp53*-/- *Rb1*-/- *Brca1-/-*, and *Trp53*-/- *Rb1*-/- *Brca2-/-* genotypes resulted in tumors histopathologically similar to HGSOC (Szabova et al. 2012). These and other models support findings that *TP53* mutations are nearly ubiquitous in ovarian carcinoma and that the *RB1* pathway is dysregulated in 67% of HGSOC tumors (Karst et al. 2011; Flesken-Nikitin et al. 2003; Chien et al. 2015; Cancer Genome Atlas Research Network 2011).

Our observation that *Pten* disruption significantly increases transformation frequency and colony size is also consistent with data from GEMMs. HGSOC-like tumor development in mouse models without *Pten* mutations have longer latencies than mice with *Pten* mutations (Zhai et al. 2017; Perets et al. 2013). Tumors with *PTEN* mutations are more aggressive and have a worse prognoses (Martins et al. 2014). *PTEN* and *RB1* mutations also significantly co-occur in HGSOC, suggesting that they may act synergistically in carcinogenesis (Gao et al. 2013; Cerami et al. 2012; Cancer Genome Atlas Research Network 2011).

In addition to identifying a core set of transformation-enhancing mutations, our data suggest that mutating two other TCGA driver genes can further enhance OSE-SC transformation susceptibility (Figure 6). Given a core set of mutations in *Trp53*, *Cdkn2a* and *Pten*, additional disruption of *Ankrd11* or *Wwox* significantly promoted adhesion-independent growth. Interestingly, genomic analyses of tumors from a mouse model deficient for *Trp53*, *Brca1*, *Brca2*, and *Pten* revealed deletions in both *Ankrd11* and *Wwox*, suggesting that they may play a role in tumor initiation or progression (Perets et al. 2013). Both *Ankrd11* and *Wwox* have also been implicated in *Trp53*-related pathways. ANKRD11 is a putative tumor suppressor that interacts with TP53 and promotes its transcription factor activity (Neilsen et al. 2008; Lim et al. 2012; Noll et al. 2012). The protein has also been shown to bind mutant TP53 and partially restore its DNA binding capacity to the *CDKN1A* promoter (Noll et al. 2012; Neilsen et al. 2008). WWOX greatly influences the response of TP53 to genotoxic stress, and *Wwox* mRNA inhibition abolishes TRP53-dependent apoptosis (Chang et al. 2001; Schrock and Huebner 2015). Mutations in *Wwox* and *Ankrd11* in our project may therefore contribute to further dysregulation of *Trp53* or may promote transformation in the presence of non-null mutations in *Trp53* induced by LentiCRISPR mutagenesis.

Several genes assessed in our study, namely *Brca2*, *Fancd2*, *Prim2*, *Brca1*, *Rad51c*, *Fat3*, and *Apc*, appeared to deter OSE-SC adhesion-independent growth when singly mutated alongside core disruption of *Trp53*, *Rb1* and/or *Pten*. However, we found that combined disruptions of subsets of these genes actually function synergistically to enhance OSE-SC transformation. Namely, co-mutation of *Brca2* and *Fancd2*, *Rad51c* and *Fat3*, or *Rad51c* and *Gabra6,* in addition to the core mutations, caused significant increases in colony growth. The growth-deterring effects of *Brca2*, *Fancd2* or *Rad51c* single mutagenesis observed in our assays have also been documented by others.

Cells lacking *Brca2* accumulate spontaneous DNA damage during G2/S phase and often senesce following the G1 checkpoint (Feng and Jasin 2017). Similarly, cells deficient in either *Rad51c* or *Fancd2* display reduced cell proliferation, especially after further induction of DNA Damage (Kuznetsov et al. 2009; Hinz et al. 2003; Kondrashova et al. 2017; Thompson et al. 2017; Tian et al. 2017). We speculate that concurrent mutagenesis of these potentially growth-deterring genes may be necessary to rescue proliferation. It remains to be investigated whether the potential synthetic lethality or growth deterrence observed here may be relevant for targeted drug development.

A fundamental aspect of our project was not only to identify and validate combinations of transformation-associated genes, but also to assess whether commonly-mutated HGSOC genes are simply passengers or unnecessary for transformation initiation. For many genes in our study, there was no evidence for involvement in OSE transformation. They include *Apc*, *Crebbp*, *Fancm*, *Nf1*, *Csmd3*, *Map2k4*, *Brca1*, and *Prim2.* Although mutations in these genes are associated with developed tumors, sequencing data alone is unable to distinguish whether mutations occurred during cancer initiation or were later events. Our study only assessed transformation initiation, so it’s possible that these genes facilitate later stages of carcinogenesis. Alternatively, many genes that are not involved in transformation initiation may be passenger mutations or a consequence of the genomic instability associated with HGSOC tumor cells.

Our study has elucidated core mutations necessary for cell autonomous transformation and has addressed fundamental questions surrounding ovarian carcinoma initiation mechanisms. However, it will be important to model the various mutation combinations *in vivo* for their abilities to induce clinically relevant cancers. Such efforts are now under way using information obtained from these studies *in vitro*. To date, TCGA has generated comprehensive genomic and transcriptomic data for 33 cancer types using over 20,000 primary tumor samples (Gao et al. 2019). These comprehensive datasets report genes that are significantly mutated or commonly deleted in different diseases, but the roles of many of these genes in cancer initiation or downstream biology is unclear. Unbiased combinatorial screening efforts, such as that we’ve applied here, in the proper cellular paradigms, may help disentangle steps of neoplastic transformation in multiple types of cancer.

## ACKNOWLEDGEMENTS

We thank Q. Sun and P. Schweitzer for providing assistance with processing and analysis of sequence data; L. Johnson for help with statistical analyses, and the staff (R. Munroe and C. Abratte) of Cornell’s transgenic facility for producing *Trp53* mutant mice. We also acknowledge technical support and advice from the following individuals: A. McNairn, L. Wang, A. Kolarzyk and M. Baccas. This work was supported by grant from the Ovarian Cancer Research Fund (#327516) and NYSTEM (C029155) to JCS and AYN, from the US National Institutes of Health (NIH) and National Cancer Institute (NCI) (CA182413) to AYN, and a predoctoral fellowship to RJY (NYSTEM C30293GG).

## METHODS

### Generation of *Trp53+/-* FVB/NJ Mice

CRISPR/Cas9 gene editing was used to generate a *Trp53+/-* co-isogenic mouse line in strain FVB/NJ. A cloning-free overlap PCR method was used to generate the DNA template for making sgRNA (Carrington et al. 2015). The following guide RNA sequence corresponding to exon 4 of *Trp53* was used: AGTGAAGCCCTCCGAGTGTC (Shalem et al. 2014; Sanjana et al. 2014). The DNA template was reverse transcribed into RNA using the MEGAshortscript T7 Transcription kit (Ambion), then purified using MinElute Columns (Qiagen; Cat#: 28004). The sgRNA (50ng/uL) and Cas9 mRNA (25ng/uL, TriLink) was microinjected into the pronuclei of FVB/NJ zygotes, then transferred injected zygotes into oviducts of pseudopregnant females. A male founder carrying a 1bp insertion in exon 4 and a predicted STOP codon before the DNA binding domain of TRP53 was selected to establish a line (Figure S1 A,B). Initial phenotyping was done after three generations, all of which were crossed to FVB/NJ animals (Figure S1 C,D).

### Mouse Embryonic Fibroblast (MEF) Isolation

MEFs were generated using previously described methodology (Todaro and Green 1963). Briefly, embryos were isolated from pregnant mice at 13 dpc (day post-coitum) and washed in PBS. Working with one embryo at a time, each embryo was placed in a clean petri dish with 0.25% trypsin EDTA solution. A small biopsy was collected for later genotyping. Sterile scalpel blades were used to mince tissue until it was able to be maneuvered with a pipette. Minced tissue incubated for 15 minutes at 37°C in trypsin solution. DMEM with 10% FBS was then used to inactivate trypsin. MEFs were plated on 10cm plates in DMEM with 10% FBS and expanded until passage 2.

### Western blotting

*Trp53*+/+, *Trp53*+/-, and *Trp53*-/- MEFs were treated with 10Gy irradiation to activate the p53 pathway. Equal quantities of cells from each group were pelleted, lysed with RIPA buffer, and utilized for immunoblotting experiments. Protein samples were collected from cell pellets lysed with RIPA buffer. Protein concentrations were normalized via both cell number and BCA assay. Samples were run through 4-15% gradient polyacrylamide gels (BioRad; Cat#: 4561083EDU) and transferred onto nitrocellulose membranes (Thermo Fisher; Cat#: 88018). We used the Rabbit anti *TRP53* primary antibody (Cell Signaling; Cat#: 9282) for detection of *TRP53*, along with goat anti-rabbit HRP-linked secondary antibody (Cell Signaling; Cat#: 7074S). Rabbit primary antibody was used for detection of *ACTB* (B-actin) (Abcam: Cat#: ab8227), along with goat anti-rabbit HRP-linked secondary antibody (Cell Signaling; Cat#: 7074S) (Figure S1C).

### LentiCRISPRv2 construct design and cloning

LentiCRISPR v2 was obtained from Addgene (plasmid # 52961; http://n2t.net/addgene:52961; RRID:Addgene 52961). We designed sgRNA guides targeting the earliest possible exon of 20 TCGA driver genes (Table 1) with the intent of inducing a gene-inactivating nonsense mutation in targets. Guides were designed using parameters described previously (Hsu et al. 2013). With few exceptions, guide sequences were chosen with an optimal off target score (>75). If no guides with a score of 75 or greater were found, we chose a guide with the highest possible score (Supplemental Table 3). Synthetic sense and antisense oligonucleotides for each guide were produced as in 25nmol quantities by Integrated DNA Technologies, such that each strand has overhangs necessary for cloning using the BSMBI restriction enzyme (New England Biolabs; Cat#: R0580S).

Target guide sequences were cloned into LentiCRISPRv2 plasmids using previously published methodology (Shalem et al. 2014; Sanjana et al. 2014). Briefly, sense and antisense oligonucleotides corresponding to each sgRNA were annealed to one another in a thermocycler (BioRad MyCycler). LentiCRISPRv2 plasmid was cut using BSMB1 restriction enzyme (New England Biolabs; Cat#: R0580S), and T4 DNA ligase (NEB; Cat#: M0202S) was used to ligate oligos into LentiCRISPRv2 plasmid. LentiCRISRv2 plasmids were transformed into One Shot Stbl3 chemically competent *E. coli* (ThermoFisher; Cat#: C737303). Transformed bacteria were plated on LB agar plates with ampicillin (Sigma-Aldrich; Cat#: A0166) for selection, and single colonies were picked and cultured in LB broth (Sigma-Aldrich; Cat#: L3147). Plasmid was isolated from bacteria using the GeneJet Plasmid Miniprep kit (ThermoFisher; Cat#: K0502) or the GeneJet Plasmid Midiprep kit (ThermoFisher; Cat#: K0481). We validated that each LentiCRISPRv2 construct contained the correct guide via Sanger sequencing (Cornell Biotechnology Resource Center) primed with the following oligonucleotide: 5’-GAGGGCCTATTTCCCATGATT-3’.

### Tissue culture of the OSN2 cell line and primary OSE

We cultured OSE, OSE-SC and OSE-NS following previously described methodology (Flesken-Nikitin et al. 2013). Briefly, ovaries were isolated from *Trp53* heterozygous FVB/NJ adult females and placed into phosphate-buffered saline (PBS) without Ca2+ or Mg2+ on ice. Ovaries were washed three times with PBS under a laminar flow hood, and a sterile scalpel blade was used to separate ovaries from the bursa.

OSE was separated from the ovary via treatment with a digestion buffer consisting of collagenase (Sigma-Aldrich; Cat#: 10269638001), dispase (Sigma-Aldrich; Cat#: 10269638001), DNaseI (Sigma-Aldrich; Cat#: 11284932001) and Bovine Serum Albumin (BSA) (Sigma-Aldrich; Cat#: A9418). We finally added the pellet from each ovary to 2mL OSE medium on gelatin-coated 24 well culture plates (Corning Costar; Cat#: CLS3527-100EA. Epithelial lineage of isolated cells was determined via detection of CK8 expression using immunofluorescent microscopy (Figure S4B-D).

Primary OSE cultures and the OSN2 cell line (Flesken-Nikitin et al. 2003) were maintained in culture in media containing DMEM (VWR; Cat#: 10-017-CM), Hams F12 (Thermo Fisher, Cat#: 11320033), 5% FBS (Atlanta Biologicals; Cat#: S11050H), hydrocortisone (Sigma-Aldrich; Cat#: H4001), insulin-transferrin-sodium selenite (Sigma Aldrich), non essential amino acids (NEAA) (Thermo Fisher; Cat#: 11140050), glutamate (Thermo Fisher; Cat#: 25030081), sodium pyruvate (Thermo Fisher; Cat#: 11360070), and penicillin-streptomycin (Thermo Fisher; Cat#: 15140122) on 0.2% gelatin-coated culture plates (Corning Costar; Cat#: 07-200-83). Cells were passaged up to two times using 0.25% Trypsin-EDTA with Phenol Red (Thermo Fisher; Cat#: 25200072) to remove adherent cells from plates.

### Immunofluorescent Microscopy

OSE, OSE-SC, OSE-NS, MEFs from FVB/N mice and OSN2 cells were grown on gelatin-coated glass cover slips for 24 hours and then fixed using methanol. Fixed cells were washed with PBS, and then blocked using goat serum (Sigma Aldrich; Cat#: NS02L). Primary Rat anti CK8 (TROMA1) antibody (University of Iowa Developmental Hybridoma Bank; Cat#: AB_531826) was added to coverslips overnight at 4°C, followed by PBS washes. Alexa Fluor 488 goat anti rat antibody (ThermoFisher; Cat#: A-11006) was added for one hour using the manufacturer-recommended concentration. One drop of mounting media (Vector Laboratories; Cat#: H-1000) was used to adhere cover slips to slides, and cells were imaged using an Olympus BX51 microscope and Olympus XM10 camera at 10x magnification. GFP expression conveyed by FUGW viral transductions was detected using a BioRad ZOE Fluorescent Cell Imager at 20x.

### Tissue Culture of HEK293T, HELA and MEF cells

HEK293T (ATCC CRL-3216) and HELA (ATCC CCL-2) cells were cultured in DMEM with 10% FBS, NEAA, and Sodium Pyruvate on gelatin-coated plates, and HELA cells were cultured in DMEM containing 10% FBS on plates coated with 0.2% gelatin. MEFs were isolated from female E13.5 embryos isolated from a breeding of two *Trp53* heterozygous parents. They were cultured on uncoated plates and were grown in DMEM containing 10% FBS. Cells were passaged onto 10cm plates using 0.25% Trypsin-EDTA with Phenol Red to remove adherent cells from plates.

### ALDEFLUOR Assay, Fluorescence-Activated Cell Sorting, and isolation of OSE subpopulations

We used the ALDEFLUOR detection kit (Stemcell Technologies; Cat#: 01700) to detect ALDH enzymatic activity in primary OSE cells. ALDEFLUOR reagent contains bodipy-aminoacetaldehyde (BAAA), which is a substrate for ALDH and can be acted upon by the enzyme. The molecule is water soluble and can pass freely into cells.

ALDH converts BAAA into bodipy-aminoacetate, which is negatively charged and consequently becomes trapped within cells with ALDH activity. Intracellular BAA accumulation leads to fluorescence. In a mixed population of cells, those with the highest levels of ALDH enzyme will convert larger quantities of BAAA to BAA, making them more fluorescent than cells with lower ALDH activity. 4-diethylamino benzaldehyde (DEAB) is an inhibitor of the ALDH enzyme and can be added to a portion of ALDEFLUOR-treated cells to act as a negative control. Inhibition of ALDH prevents high levels of BAAA conversion to BAA, resulting in lower levels of fluorescence.

Cells (4x106) were placed in ALDEFLUOR buffer and active reagent according to the manufacturer’s protocol. A subpopulation of ALDEFLUOR-treated cells was also treated with DEAB as a negative control. Fluorescence activated cell sorting (FACS) of ALDEFLUOR-treated cells was performed on an Aria II sorter using FACS DiVa software (BD Biosciences). The brightest 2–5% of ALDEFLUOR-treated cells were identified and gated electronically based on their characteristic light-scatter properties on the fluorescein isothiocyanate (FITC)-channel emission pattern after excitation with a 13–20 mW, 488-nm ellipse-shaped laser (elliptical) BD FACSAria II. The ALDH fluorescence emissions were captured simultaneously through a 515/20-nm band-pass and 505-nm long-pass filter. ALDH+ (OSE-SC) and ALDH− (OSE-NS) OSE cells were collected in 5ml falcon tubes, cultured, and were subjected to lentiviral transduction and colony formation assays (Figure S4A).

### Viral Packaging

The following vectors were used for lentovirus packaging: PsPax2 (Addgene plasmid #12260); VSV-G (Addgene plasmid #8454); LentiCRISPRv2 (Addgene plasmid #52961). HEK293T cells were transfected with 10ug LentiCRISPRv2, 7.5ug PsPax2 and 2.5ug VSV-G using TransIT-LT1 transfection reagent via manufacturer instructions (Mirus; Cat#: MIR 2305). FUGW (Addgene; Cat#: 14883), a GFP-expression lentiviral construct, was also packaged separately to perform control viral transduction experiments (Figure S5, S6, S7).

Following transfection of HEK293T cells, cell media was collected after 48 hours and 72 hours. Media was concentrated via centrifugation in Amicon Ultra-15 columns (Millipore; Cat#: UFC903024) such that final volume was 500uL. Concentrated virus was filtered using 0.45um syringe filters (ThermoFisher; Cat#: 725-2545) and immediately added to cultured OSE in 24-well plates. Cells were transduced with lentivirus for 48 hours. After transduction, viral media was replaced with OSE media.

### Viral Titer Calculation

LentiCRISPRv2 constructs have a puromycin resistance gene, which allows any cell transduced with a LentiCRISPRv2 virus to survive puromycin treatment. Puromycin survival amongst a population of transduced cells is therefore a function of higher viral MOI. Higher survival due to high MOI is also negatively correlated with single infection percentage, since a higher number of viral particles in solution increases the probability that a given cell will receive more than one transduction. The mathematical relationship between puromycin survival and MOI/SIP has been described by Sanjana et al. 2014 (Sanjana et al. 2014) (Figure S8A). If no minilibrary genes influence cell growth, then any gene could be expected to be targeted at a random rate. The random rate of gene targeting is equal to the number of ways a cell could receive at least one of three LentiCRISPRs targeting a specific gene from 60 total, divided by the total number of possibilities. Because there are 7 functional viral particles per cell, the total number of ways to get one of 3 LentiCRISPRs is 607. The number of ways that a different gene can be targeted is 577. The total number of possibilities is 607. We therefore determined that the rate of random gene targeting given a library with MOI of 7 is 30.2% (Figure S8B).

### Minilibrary functional validation using next-generation sequencing

OSN2 cells (*Trp53*+/-) were transduced with all 60 minilibrary constructs at a MOI of 7 for 48 hours, and all transduced cells were collected following brief culture. Cells were spun down, then genomic DNA was isolated from cells using the Agencourt DNAdvance DNA isolation kit (Beckman Coulter; Cat#: A48705). All LentiCRISPR target sites (except for those targeting *Trp53*) were amplified using PCR, and amplicons were barcoded (Supplementary Table 1). Successful amplification of all LentiCRISPRv2 target sites was verified via agarose gel electrophoresis to assess amplicon size. We also performed Sanger sequencing on all PCR products to confirm that all intended regions were amplified. Following verification, all reactions were pooled into a single tube and purified using the QiAquick PCR purification kit (Qiagen; Cat#: 28104) (Figure S2A). LentiCRISPRs targeting *Trp53* were excluded from initial verification experiments because OSN2 cells lack *Trp53* alleles but were later assessed using a Surveyor mutagenesis assay (Integrated DNA Technologies; Cat#: 706020).

300bp paired end sequencing was performed using Illumina MiSeq to detect indels in minilibrary target site amplicons at a read depth of 25 million reads. BWA MEM software was used for genome alignment (arXiv:1303.3997). Insertions or deletions greater than 4 base pairs in all minilibrary target sites were then tallied in transduced and untransduced control cells. Individual minilibrary constructs were considered “functional” if two-fold more indels were present in transduced cells compared to untransduced cells. We found that most constructs were functional (Figure S2B). Non-functional LentiCRISPRs (less than two fold difference in number of indels) or those targeting *Trp53* were redesigned and functionally validated using a Surveyor mutagenesis assay (Figure S3).

### Surveyor Mutagenesis Assay

The Surveyor mutagenesis assay was used to validate activity of some vectors (Figure S3). OSE cells were transduced with *Trp53*-targeting LentiCRISPRs because OSN2 cells lack *Trp53* alleles. OSN2 cells were transduced with redesigned LentiCRISPRs intended to replace non-functional constructs. Briefly, LentiCRISPR target sites were PCR-amplified in transduced (edited) and untransduced (control) OSE cells (Supplemental Table 2). Amplicons from control and edited cells were mixed in equal concentrations, heated to cause separation of complementary strands, then cooled to cause re-annealing of complementary strands. If a mutation is present in the transduced cell amplicons, then heteroduplexes containing several unmatched basepairs will form as amplicons from LentiCRISPR-transduced cells try to anneal with amplicons from untransduced cells. As a control, amplicons from untransduced cells were mixed with amplicons from other untransduced cells. Heating and re-annealing of amplicons from untransduced cells are not expected to cause mismatches in re-annealed DNA, since amplicons do not contain LentiCRISPR-induced mutations and should all be identical. Surveyor nuclease recognizes mismatches in annealed amplicons and cleaves DNA at that site. Therefore, if a LentiCRISPR-induced mutation(s) is present in amplicons from transduced cells, and DNA from those amplicons are annealed to non-mutated amplicons from the same genomic region, a mismatch would occur and the site would be cut by Surveyor nuclease. Surveyor nuclease was added to both experimental and control groups, and all samples were run through a 2% agarose gel. Bands unique to transduced samples indicate that mutagenesis of transduced cell target sites has occurred. Numbers displayed below gene names refers to minilibrary construct ID number. We observed unique bands in all transduced DNA samples, suggesting that all replacement LentiCRISPRs and *Trp53*-targeting LentiCRISPRs possess editing ability.

Efforts to functionally assess all minilibrary constructs resulted in a validated minilibrary of 60 constructs, including one construct targeting *Gfp* as a negative control (Figure S2, S3). *Rad51c* was only targeted by two LentiCRISPRs in the finalized library due to failed validation of a third construct.

### Soft Agar Culture, Colony Isolation, and Imaging

Adhesion independent growth was assessed using Cell Biolabs Inc 96 Well Cell Transformation Assay (Cell Biolabs; Cat#: CBA-135). Cells were plated at a density of 3000 cells per well in a 48 well dish and were suspended in 150ul agar/media solution. Transformation was monitored for one week, and colonies were collected via the manufacturer’s instructions. In short, Cell Biolabs Inc matrix solubilization solution was added to each well of cells (Cell Biolabs; Cat#: CBA-135). Colonies were resuspended in OSE media and plated at very low density on 15cm plates. Individual, distinct colonies growing on 15cm plates were picked and cultured independently. Sterile filter paper was soaked in trypsin and was used to pick individual colonies from plates.

Picked colonies were added to 24-well plates and were passaged using OSE culturing methodology. Brightfield images of culture wells were taken using a Nikon SMZ1500 microscope and Nikon Digital Sight DS-Fi1 camera system at 2x magnification. 10x and 20x images were taken using a Nikon TMS-F microscope and Moticam 2300 3.0MP Live Resolution camera system.

### LentiCRISPRv2 identification via Next Generation Sequencing

Individual transformants from soft agar assays were isolated and expanded in culture as adherent cell lines. 500,000 cells per transformant were grown, spun down, and lysed for DNA extraction using the Agencourt DNAdvance DNA isolation kit. We designed a single pair of primers flanking unique LentiCRISPR guide sequences to amplify all genome-integrated constructs in each sample. We used the following primers: CTTGGCTTTATATATCTTGTGGAAAGG and CGACTCGGTGCCACTTT.

Illumina overhangs were also added to each primer. PCR reactions were performed on genomic DNA isolates from each individual colony, resulting in amplification of any genome-integrated LentiCRISPRv2 construct. Amplicons corresponding to individual transformants were uniquely barcoded and library prep was completed using the Miseq Reagent Kit v2 (illumina; Cat#: MS-102-2001) according to the manufacturer’s instructions. 2 x 251bp paired end sequencing was performed on pooled, uniquely barcoded amplicons.

We created a custom “genome” of guide sequences and used BWA MEM to align reads from uniquely barcoded transformants to guide sequences. Each alignment to a particular guide represents a “hit”, meaning that a particular genome-integrated guide was PCR-amplified in transformant DNA isolates, and was detected via sequencing.

We calculated the average number of reads per construct and determined the average background read count. We designated any individual construct as genome-integrated if it had more aligned reads than twice the background read count. We finally performed hierarchical clustering of integration data to best visualize patterns of LentiCRISPR integration.

### Mice and Genotyping

All animal use was conducted under protocol (2004-0038) to J.C.S. and approved by Cornell University’s Institutional Animal Use and Care Committee.

We followed previously described methodology to generate crude tissue lysates for PCR (Truett et al. 2000). The following forward and reverse primers were used for PCR and genotyping: TTGTTTTCCAGACTTCCTCCA and GCATTGAAAGGTCACACGAA. PCR success was confirmed on an agarose gel, and the forward primer was re-used for Sanger sequencing.

### Materials Availability

All LentiCRISPRv2 plasmids, *Trp53+/-* FVB/NJ mice, and all transformed cell lines, generated in this study are available upon request.

