## Supplemental Figures and Tables for "Most commonly mutated genes in High Grade Serous Ovarian Carcinoma are nonessential for ovarian surface epithelial stem cell transformation"

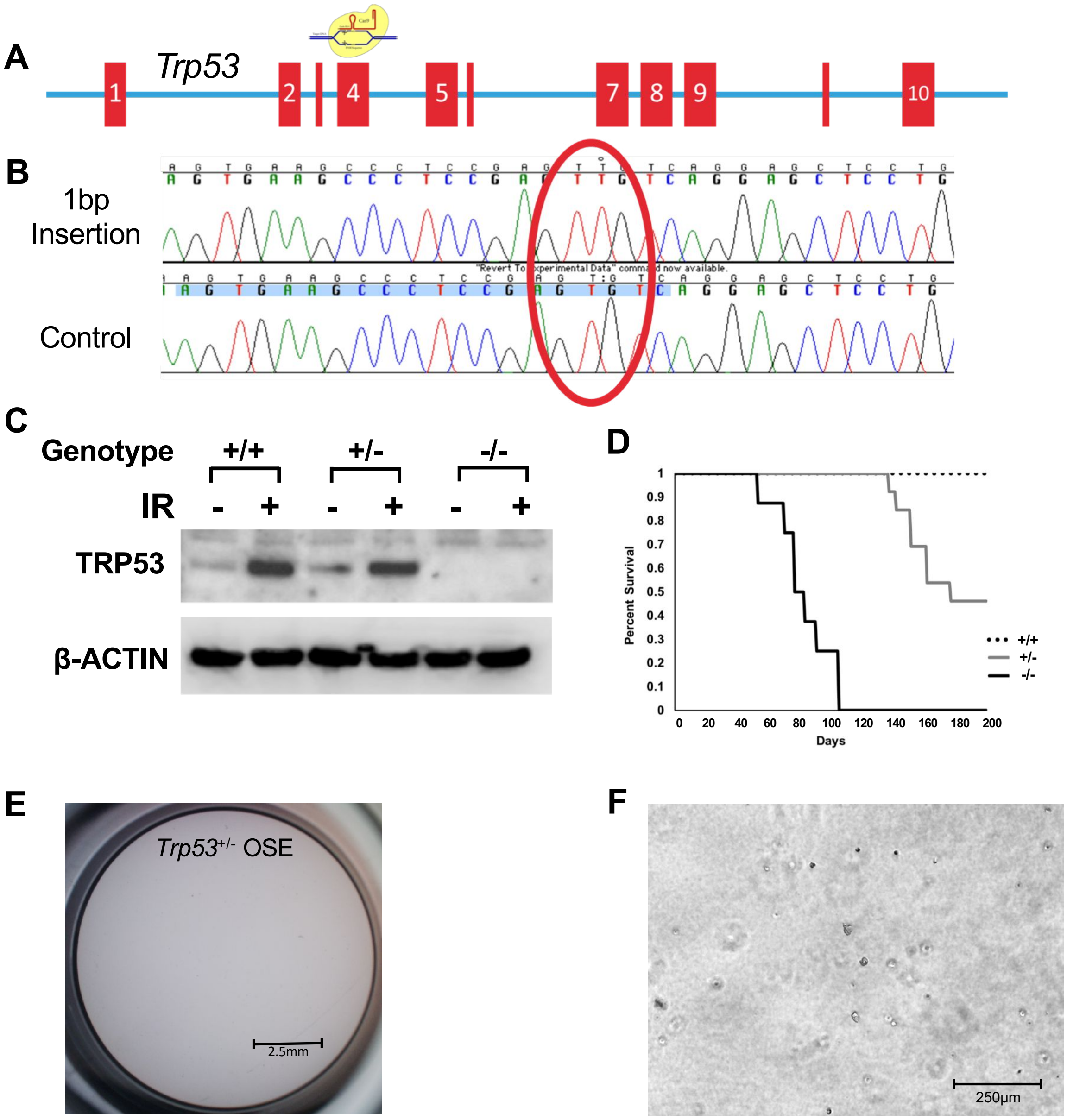


**Supplemental Figure 1: Generation of a *Trp53* null mouse line congenic in strain FVB/NJ by genome editing and validation. (A)** Schematic of mouse *Trp53* gene and CRISPR/Cas9 targeting of exon 4, which precedes the DNA binding domain of TRP53. **(B)** Sanger sequencing chromatogram of the founder animal used to establish the line. The CRISPR/Cas9 target site is highlighted in blue, and the 1bp insertion in circled in red. **(C)** Western blot analysis of TRP53 levels in MEFs treated (or not) with 10 Gy of ionizing irradiation (IR). Increased TP53 (due to stabilization) was evident in WT and heterozygous MEFs, but the protein was completely absent in homozygous mutants. ACTB (β-actin) was also detected as a control for equal loading. **(D)** Kaplan-Meier survival plots of female *Trp53*$^{+/+}$, *Trp53*$^{+/-}$, and *Trp53*$^{-/-}$. **(E,F)** *Trp53*$^{+/-}$ ovarian surface epithelium (OSE) cells do not transform in the soft agar assay. No adhesion independent growth was noted for *Trp53*$^{+/-}$ OSE after 14 days.

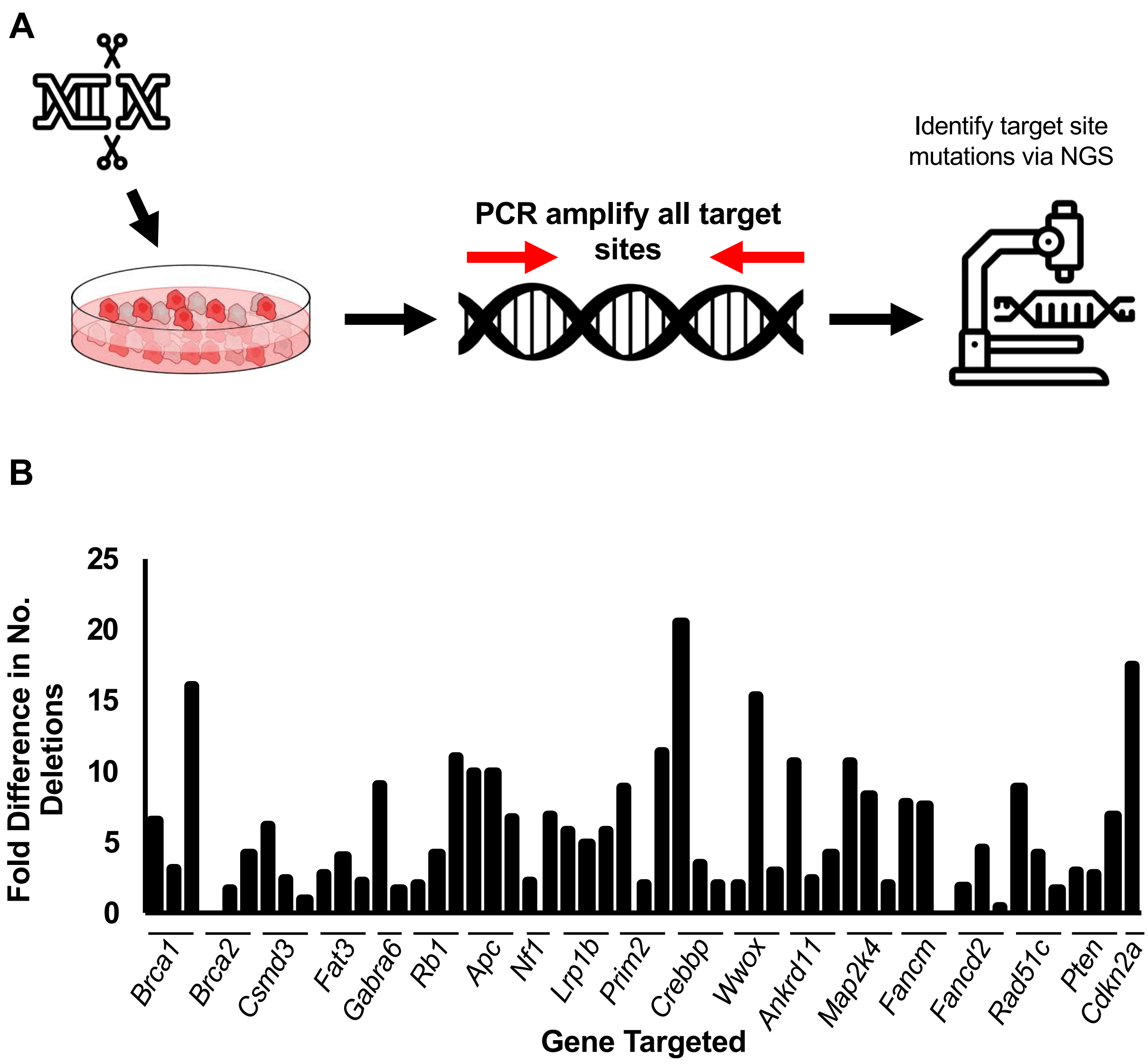


**Supplemental Figure 2. Next generation sequencing of minilibrary LentiCRISPRv2 target sites suggests that most constructs cause indels at target sites. (A)** Schematic representation of minilibrary validation strategy. **(B)** Fold difference in number of 4+ base pair indels at minilibrary target sites between minilibrary-transduced OSN2 and untransduced OSN2. All minilibrary target sites were PCR-amplified and sequenced using NGS. Each gene name on the X axis represents an individual LentiCRISPRv2 target site. All target sites were assessed except for *Trp53* because OSN2 cells lack *Trp53* alleles. All three *Cdkn2a* target sites are contained within the same amplicon. Most minilibrary target sites in transduced cells exhibited two-fold or more indels vs target sites in untransduced cells. Constructs unable to cause two-fold or more deletions in target sites vs controls were considered "non-functional" and were redesigned. Target sites that were not amplified or not sequenced were also later validated or redesigned. (NGS = Next Generation Sequencing)

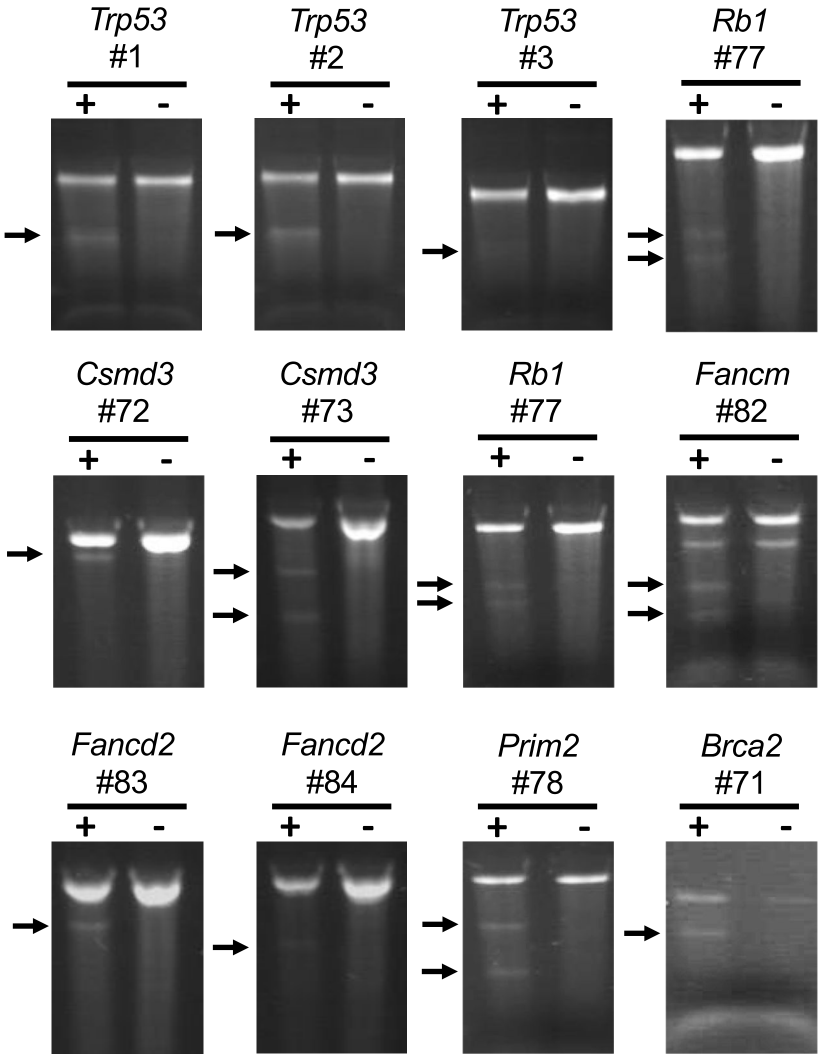


**Supplementary Figure 3. Redesigned LentiCRISPRs and LentiCRISPRs targeting Trp53 cause mutations in target sites.** LentiCRISPRv2 target sites were PCR-amplified, and mutations were detected using the Surveyor mutagenesis assay (See Methods). The "+" lane represents Surveyor-treated heteroduplexes between edited target site amplicons and corresponding unedited target site amplicons. The "-" lane contains Surveyor-treated re-annealed amplicons from control, unedited cell DNA. Bands in the "+" lane suggest that a LentiCRISPRv2-induced mutation was present within the edited amplicon, resulting in mismatched DNA and digestion by Surveyor nuclease.  Bands in the "-" lane represent background digestion by Surveyor.  Arrows indicate bands unique to edited samples, suggesting that mutagenesis following LentiCRISPRv2 targeting occurred.

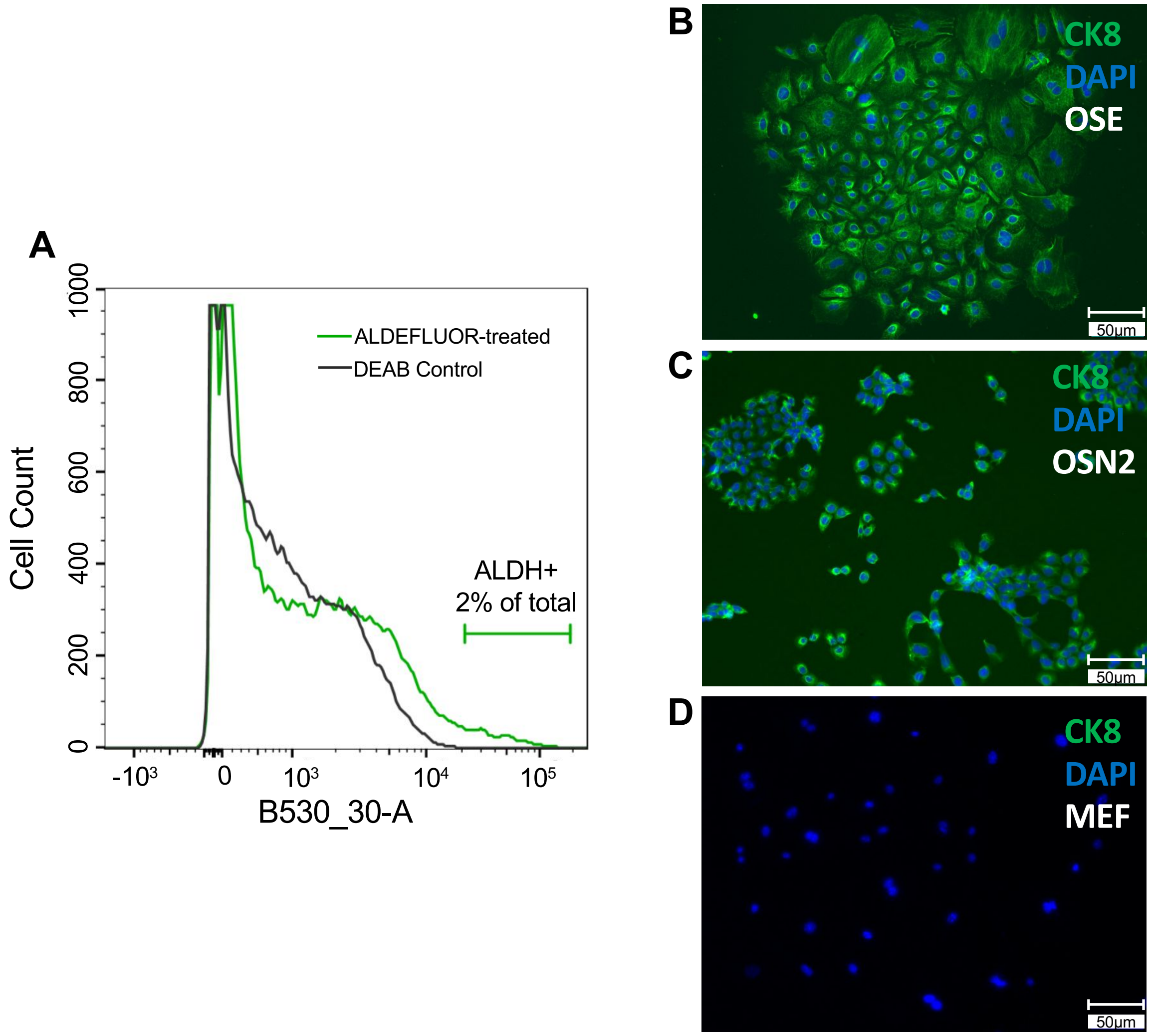


**Supplemental Figure 4: OSE contains a small population of ALDH+ cells and express the CK8 epithelial marker in culture.** (**A)** Isolation of OSE-SC and OSE-NS via FACS sorting. OSE cells were treated with ALDEFLUOR reagent (green line) or ALDEFLUOR plus the DEAB ALDH inhibitor (grey line) (see Methods). ALEFLUOR-treated cells that fluoresced more than DEAB control cells were isolated as an OSE-SC enriched population. Cells with low levels of fluorescence were isolated as OSE-NS. **(B-D)** CK8 epithelial cell marker expression in OSE. OSE was isolated from *Trp53*$^{+/-}$ FVB/N mice. OSN2 cells are an ovarian surface epithelial cell line and a positive control for CK8 expression, and MEFs from *Trp53*$^{+/-}$ FVB/NJ mice were chosen as a negative control. CK8 was expressed in OSN2 cells and OSE, but not in MEFs.

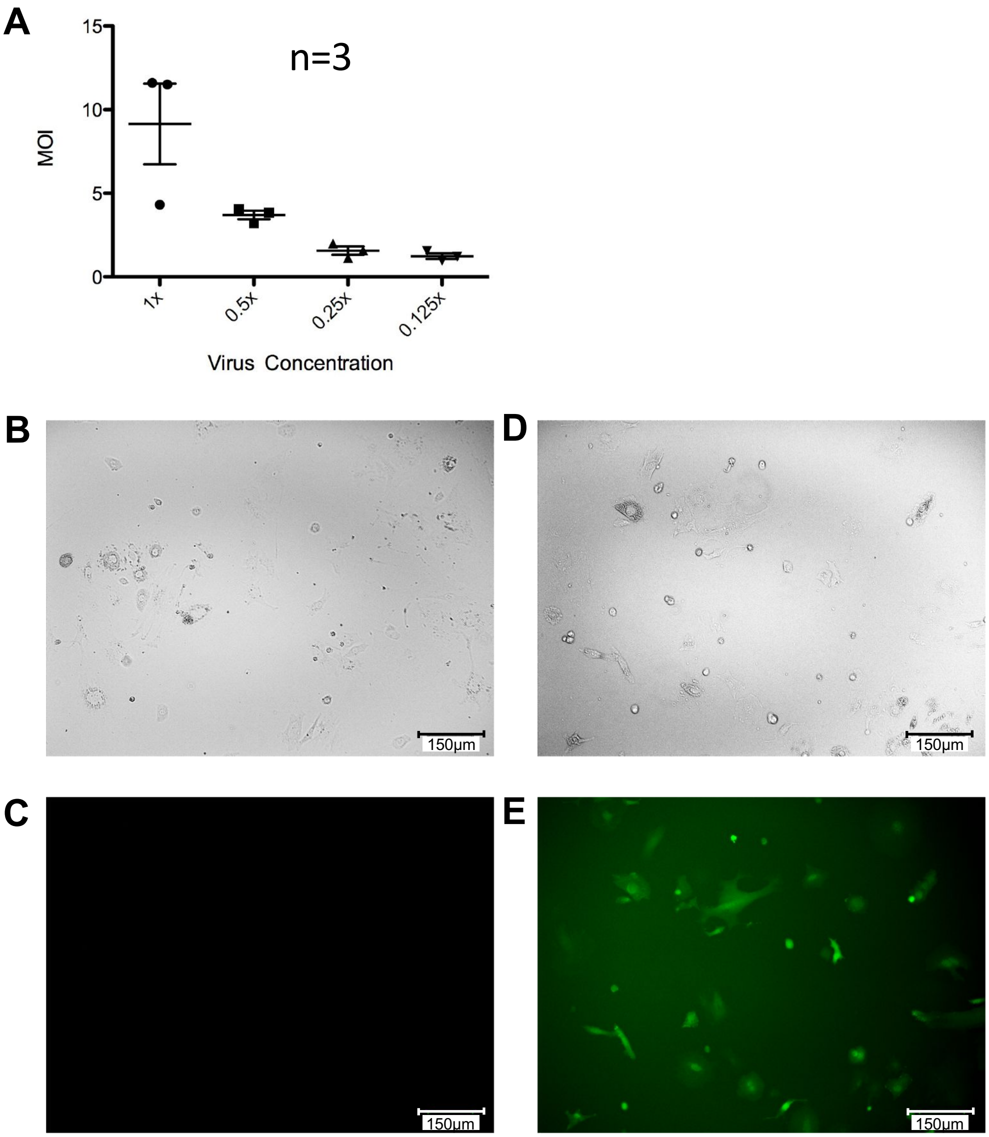


**Supplemental Figure 5. Assessment of minilibrary viral titer and FUGW (GFP-expressing lentiviral construct) transduction efficiency. (A)** LentiCRISPRv2 MOI in response to serial dilution of concentrated lentivirus. **(B-E)** Expression of GFP in OSE cells transduced with FUGW (GFP-expressing) lentivirus. OSE cells were transduced with FUGW **(D,E)** or remained Untransduced as a control **(B,C).** FUGW-transduced OSE **(D,E)** ubiquitously expressed GFP. No fluorescence was observed for untransduced OSE following imaging using the same exposure time **(B,C).**

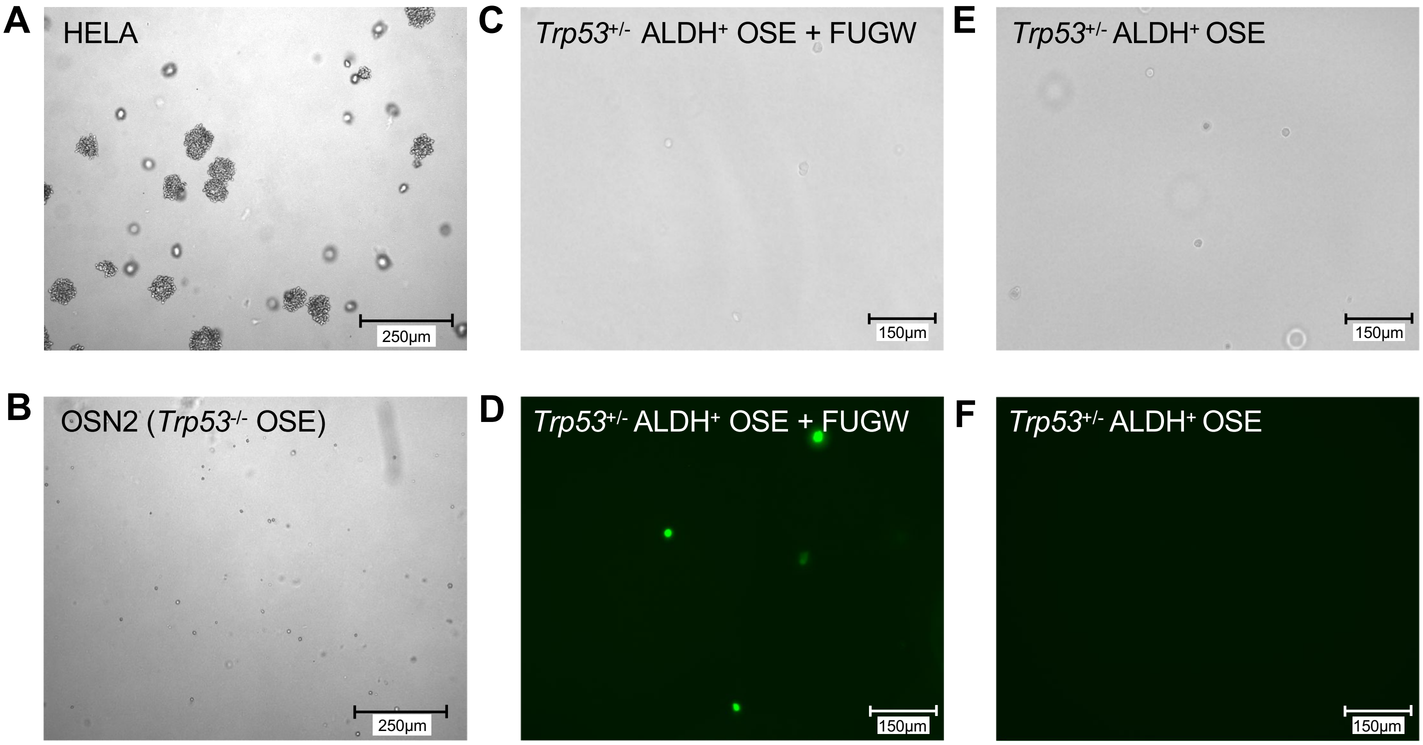


**Supplemental Figure 6: *Trp53* mutations and lentiviral transduction are not sufficient for adhesion independent growth of OSE.** HELA cells **(A)**, *Trp53*$^{+/-}$ OSE-SC **(E,F)**, *Trp53*$^{-/-}$ OSE (OSN2) **(B)**, and FUGW-transduced (GFP-expressing lentivirus) *Trp53*$^{-/-}$ OSE-SC **(C,D)** were plated in soft agar. Colony growth was noted for HELA cells **(A)**, but not for any OSE genotype or treatment group **(B-F)**. No colonies were observed following FUGW transduction, but GFP expression was noted **(C,D)**. No GFP expression was observed for untransduced *Trp53*$^{+/-}$ OSE-SC **(F)**.

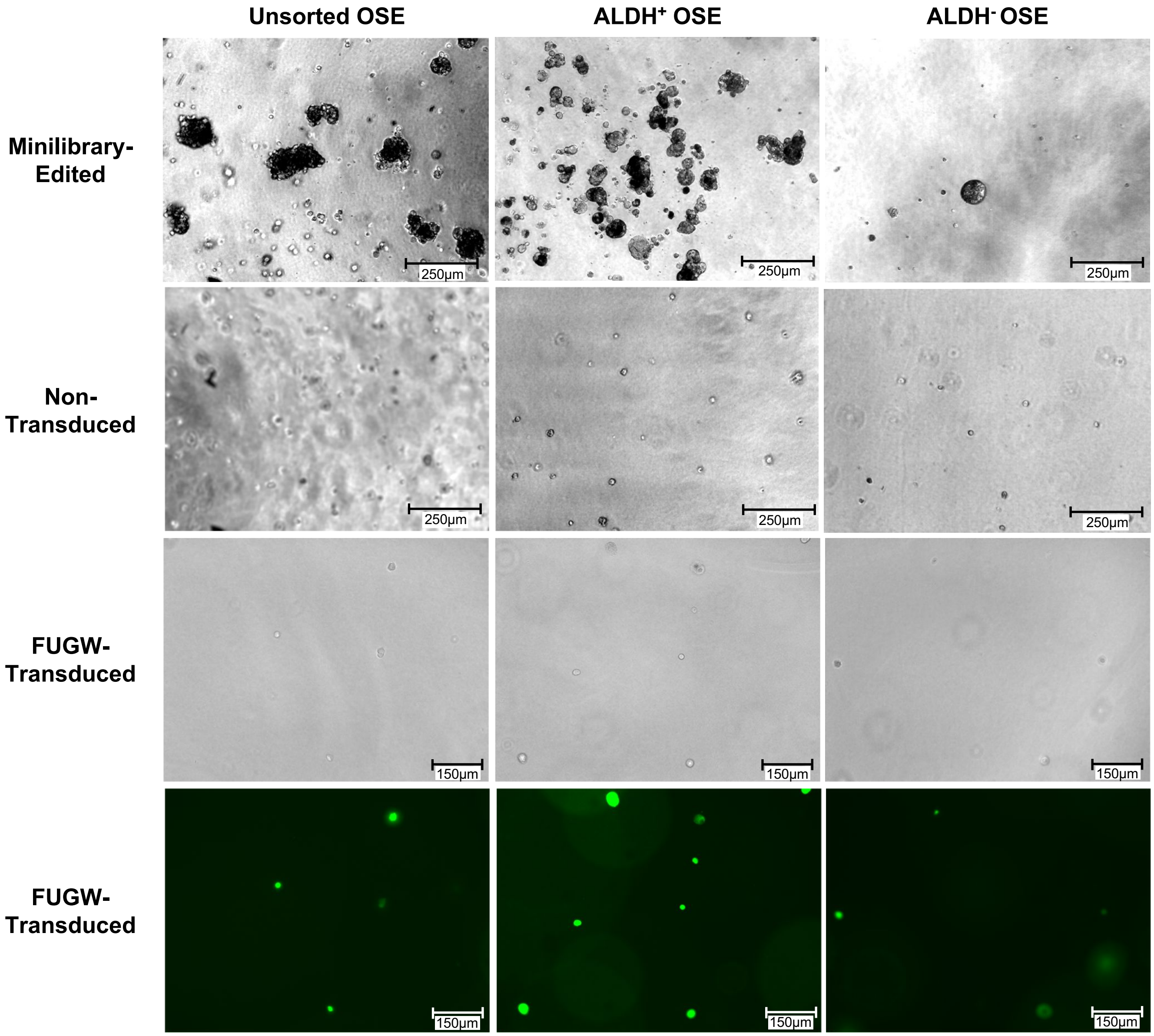


**Supplemental Figure 7. Minilibrary transduction, but not FUGW (GFP-expressing lentivirus) transduction, causes transformation of *Trp53*$^{+/-}$ unsorted OSE, OSE-SC, and OSE-NS.** The x axis indicates the OSE population being treated. The y axis indicates the treatment given to a corresponding OSE population. Only minilibrary virus transduction resulted in colony growth for all groups. FUGW transduction resulted in GFP expression but no adhesion independent growth. Cells also do not transform if untransduced.

**A**

### Determination of Single Infection Percentage (SIP), n Viral Particles, and Multiplicity of Infection (MOI):

SIP and MOI were calculated using the Poisson Distribution.
m = MOI
n = Number of viral particles
P(survival) = puromycin survival

$$P(n) = \frac{m^n e^{-m}}{n!}$$

$$p(survival) = P(n > 0) = 1 - P(n = 0) = 1 - e^{-m}$$

Solving for SIP as a function of survival:

$$SIP = \frac{P(n=1)}{P(n \geq 1)} = \frac{P(n=1)}{P(n>0)} = \frac{\big(1 - p(survival)\big)\ln(1 - p(survival))}{p(survival)}$$

**B**

### Probability of a single gene being targeted at random:

$$\frac{Ways\ to\ get\ at\ least\ 1\ of\ 3\ LentiCRISPRs from\ 60\ total}{Total\ Number\ of\ Possibilities} = \frac{60^7 - 57^7}{60^7} = 0.302$$

### Probability of GFP being targeted in any cell:

$$\frac{\#\ ways\ to\ get\ 1\ of\ 60\ LentiCRISPRs}{Total\ Number\ of\ Possibilities} = \frac{60^7 - 59^7}{60^7} = 0.111$$

**Supplemental Figure 8. Calculation of MOI (multiplicity of infection), SIP (single infection percentage) and random gene targeting frequency**. **(A)** Determining MOI and SIP as a function of LentiCRISPR-transduced cell puromycin survival. **(B)** Probability of random gene targeting in a population of cells transduced with virus with a MOI of 7.

**A**

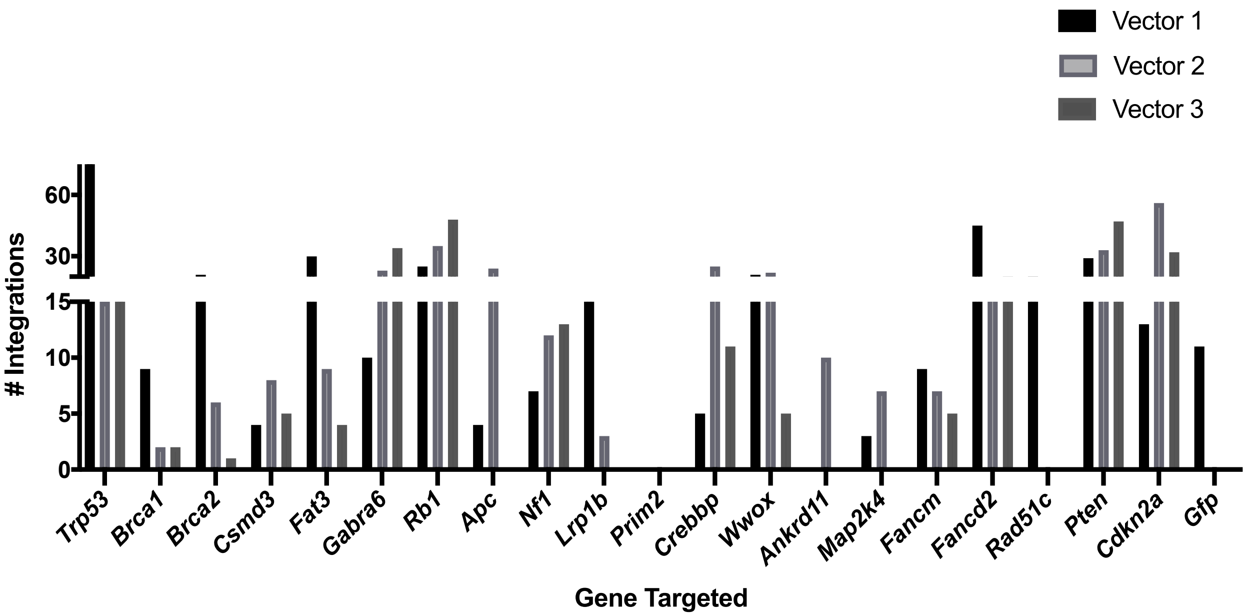


**B**

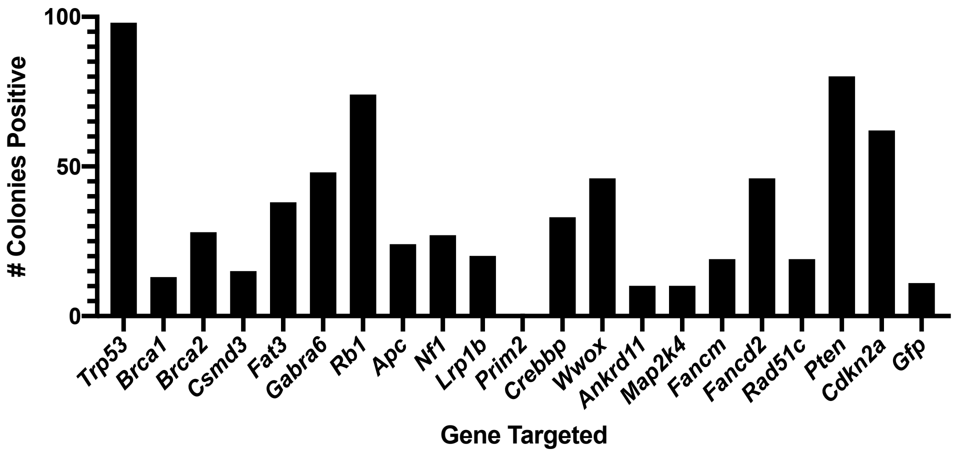


**Supplemental Figure 9: Frequently gene "hits" in OSE-SC transformants were targeted by three unique LentiCRISPRv2 constructs. (A)** Quantification of individual LentiCRISPRv2 construct genome integrations in OSE-SC transformants. Each bar corresponding to a target gene (x axis) represents an individual LentiCRISPRv2 construct. All genes that were targeted in 30% or more of OSE-SC samples had representation from all three gene-targeting LentiCRISPR constructs. **(B)** Gene targeting frequency among all OSE-SC transformants. LentiCRISPRv2 constructs targeting each putative HGSOC driver gene were identified in OSE-SC colonies using next generation sequencing. The frequency by which individual genes were targeted by at least one LentiCRISPRv2 construct among all OSE-SC transformants was tallied.

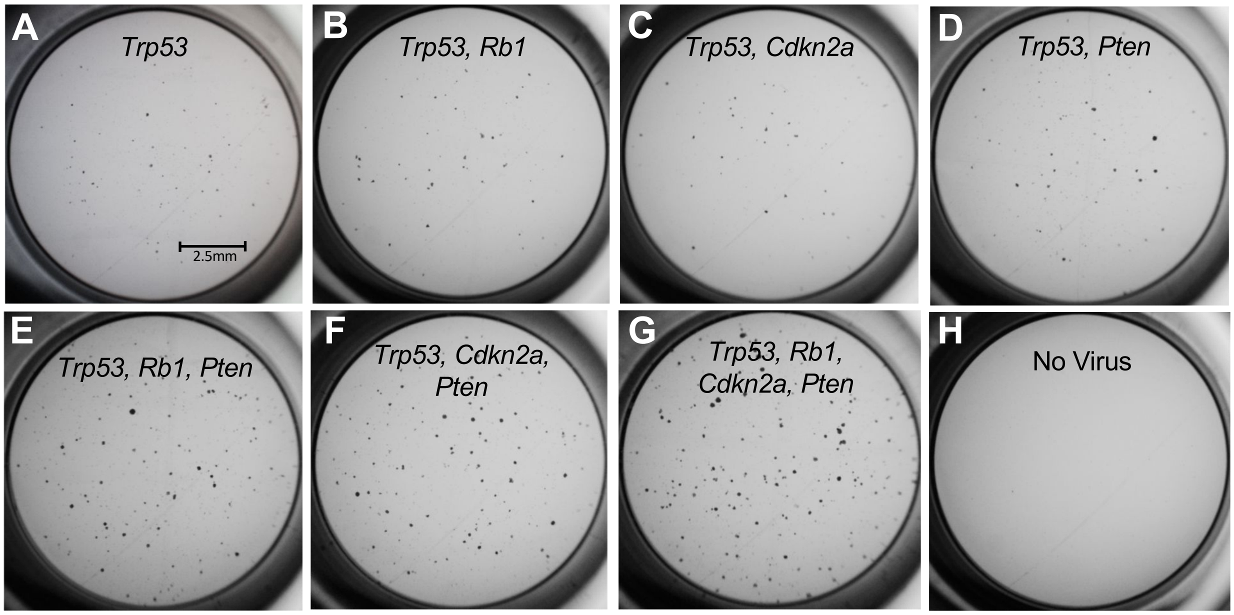


**Supplementary Figure 10.** ***Trp53, Rb1, Cdkn2a, and Pten*** **function synergistically in OSE-SC transformation initiation.** Cells were transduced with LentiCRISPRs targeting *Trp53* alone **(A)**, *Trp53* and *Rb1* **(B)**, *Trp53* and *Cdkn2a* **(C)**, *Trp53* and *Pten* **(D)**, *Trp53*, *Rb1* and *Pten* **(E)**, *Trp53*, *Cdkn2a* and *Pten* **(F)**, *Trp53*, *Rb1*, *Cdkn2a*, and *Pten* **(G)**, or were not transduced as a negative control **(H)**. Adhesion independent growth was observed for all groups except for the untransduced control. The quantity of transformants in each group was tallied for each replicate (n=6). Significant increases in adhesion independent growth vs *Trp53* alone were noted for groups **E, F** and **G** (Students t-test, $p<0.05$).

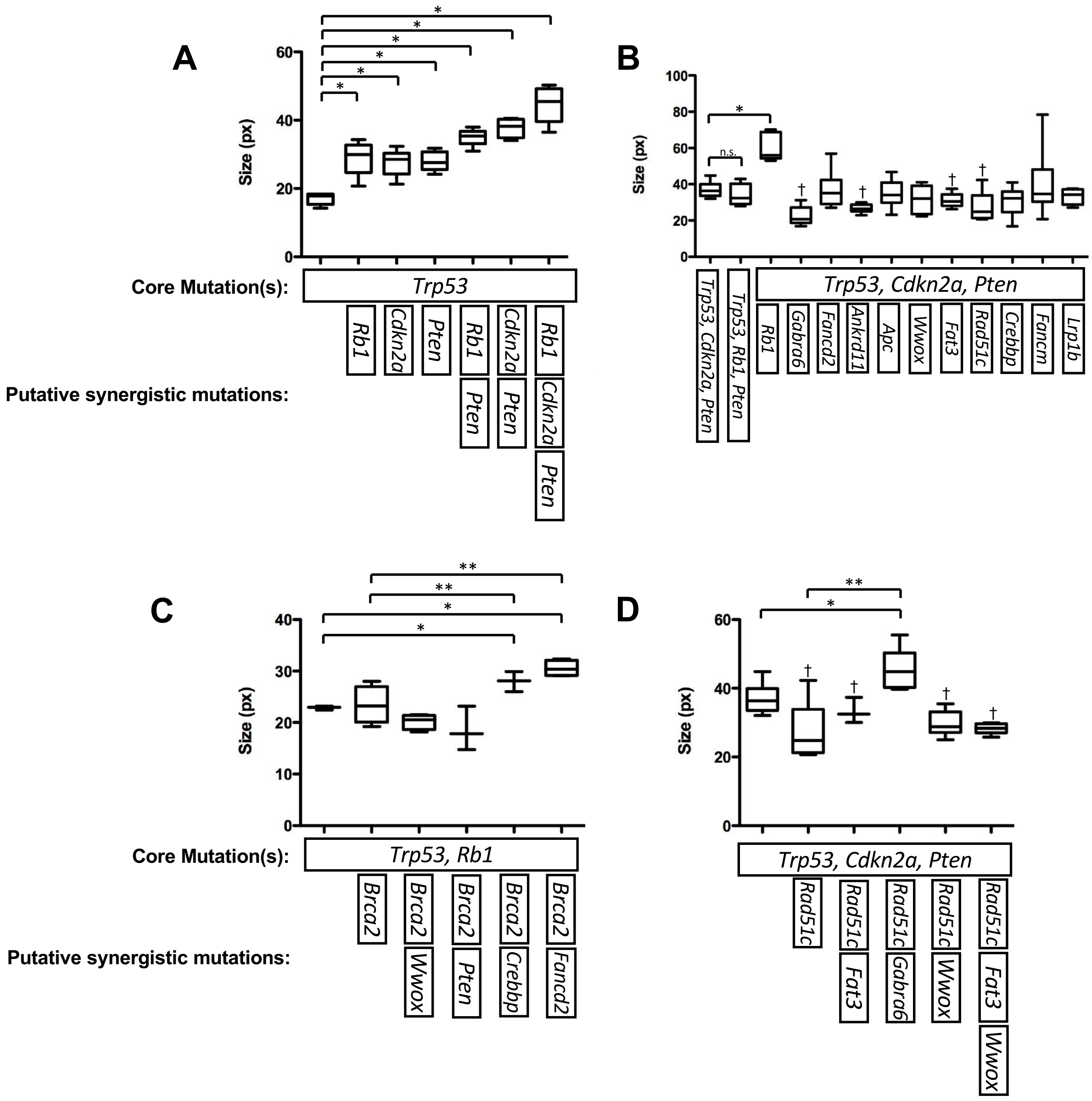


**Supplemental Figure 11. Colony size resulting from OSE-SC transformation with targeted sets of LentiCRISPRv2 constructs.** A baseline level of adhesion independent growth was first assessed via induction of specific "core mutations". Additional minilibrary target genes were then mutated alongside core mutations to assess whether they act synergistically to promote adhesion independent growth. **(A)** Targeted transduction of *Trp53*, *Rb1*, *Pten* and *Cdkn2a* LentiCRISPRs in OSE-SC. (n=6). **(B)** Targeted transduction of *Trp53*, *Cdkn2a* and *Pten* LentiCRISPRs plus putative transformation enhancers. **(C)** Targeted transduction of *Brca2* LentiCRISPRs and *Brca2*-associated LentiCRISPRs. **(D)** Targeted mutagenesis of *Rad51c* LentiCRISPRs and *Rad51c*-associated LentiCRISPRs. Values higher than baseline with Students' two-tailed t-test $p<0.05$ are labeled with an asterisk (*), and those lower than baseline are labeled with an obelisk (†). Standard error of the mean (SEM) error bars. N = 6 in all cases.

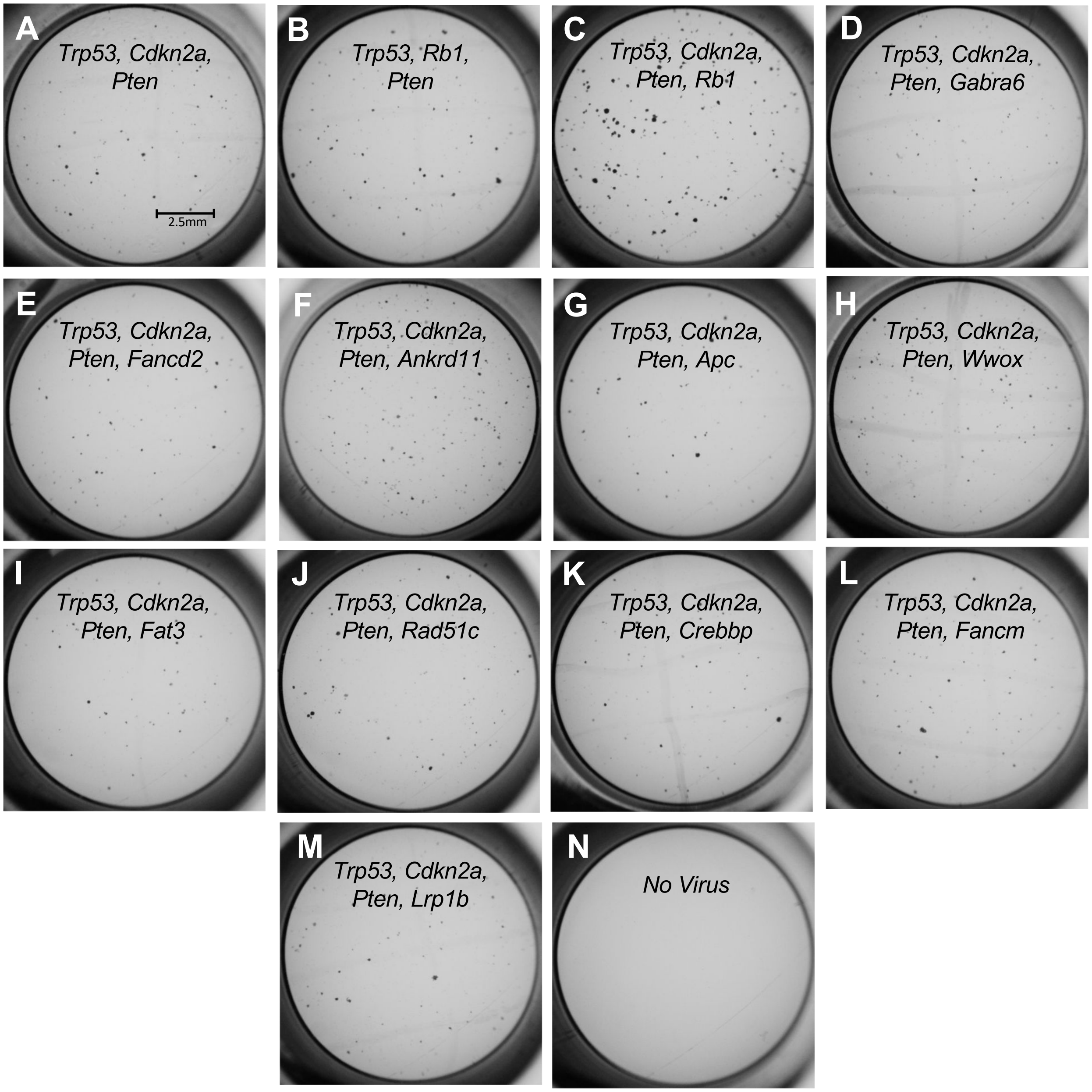


**Supplemental Figure 12: LentiCRISPRv2 targeting of *Ankrd11* or *Wwox* alongside *Trp53, Cdkn2a*, and *Pten* causes significant increases in adhesion independent growth vs combined mutagenesis of *Trp53, Cdkn2a*, and *Pten*.** Cells were transduced with core mutations in *Trp53*, *Cdkn2a* and *Pten* plus *Rb1* **(C)**, *Gabra6* **(D)**, *Fancd2* **(E)**, *Ankrd11* **(F)**, *Apc* **(G)**, *Wwox* **(H)**, *Fat3* **(I)**, *Rad51c* **(J)**, *Crebbp* **(K)**, *Fancm* **(L)**, *Lrp1b* **(M)**, or were untransduced **(N)**. Adhesion independent growth was noted for all groups except for the untransduced control. The quantity of transformants in each group was tallied for each replicate (n=6). Significant increases in adhesion independent growth were observed following the addition of LentiCRISPRs targeting *Rb1*, *Wwox* or *Ankrd11* (Students t-test, $p<0.05$). Significant decreases in adhesion independent growth were noted following the addition of Fancd2, Apc, Fat3, or Rad51c (Students t-test, $p<0.05$).

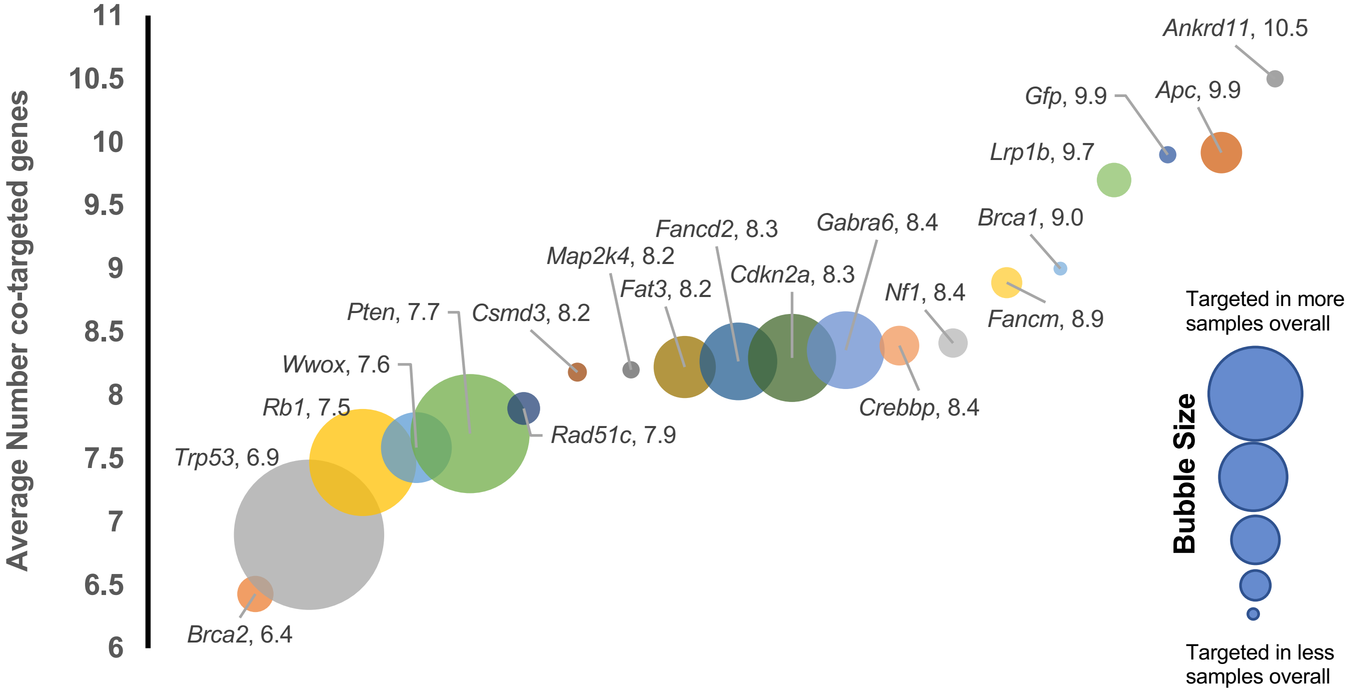


**Supplemental Figure 13. Underrepresented minilibrary target genes are frequently found in colonies with greater overall quantities of targeted genes in OSE-SC.** The Y axis represents the average number of genes targeted alongside the corresponding gene in the X axis. Bubble size corresponds to the overall number of samples that contain LentiCRISPRs targeting a given gene. Smaller bubbles therefore indicate that a gene is targeted infrequently in OSE-SC samples overall, while a larger bubble indicates that a given gene is frequently targeted. Genes targeted in few samples overall correlate positively with the number of concurrently targeted genes.

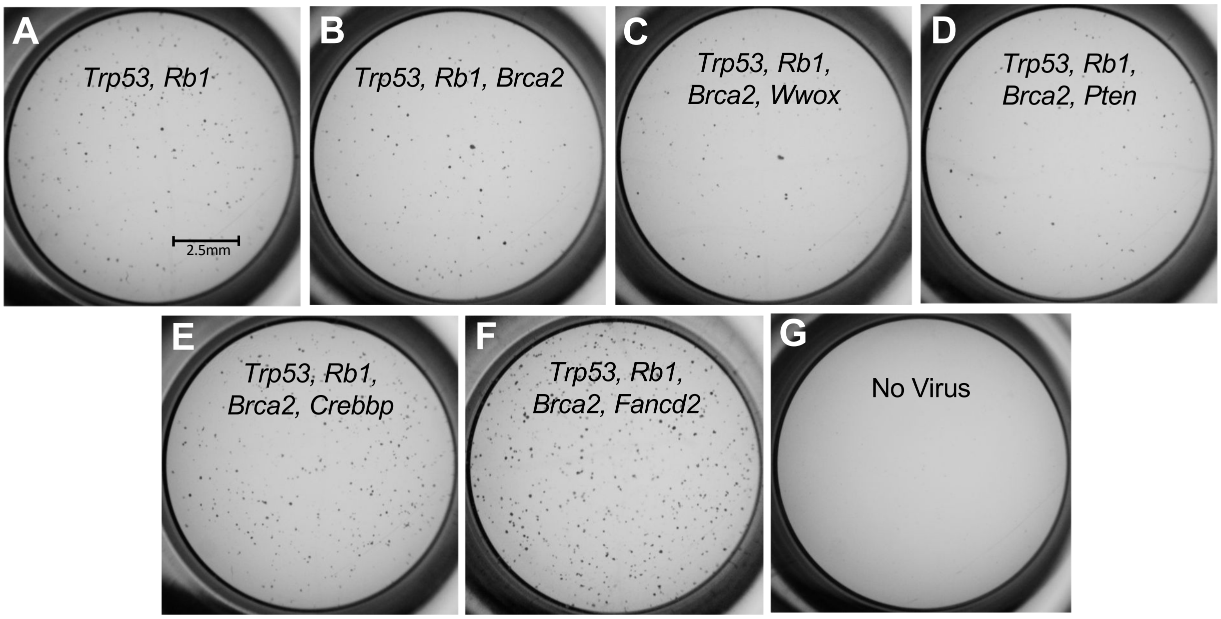


**Supplemental Figure 14. *Fancd2* and *Brca2* function synergistically to promote OSE-SC adhesion independent growth.** Cells were transduced with LentiCRISPRs targeting *Trp53* and *Rb1* **(A)**, *Trp53*, *Rb1* and *Brca2* **(B)**, *Trp53*, *Rb1*, *Brca2* and *Wwox* **(C)**, *Trp53*, *Rb1*, *Brca2* and *Pten* **(D)**, *Trp53*, *Rb1*, *Brca2* and *Crebbp* **(E)**, *Trp53*, *Rb1*, *Brca2* and *Fancd2* **(F)**, or were not transduced as a negative control **(G).** Adhesion independent growth was observed for all groups except for the untransduced control. The quantity of transformants in each group was tallied for each replicate (n=5). Significant increases in adhesion independent growth vs combined mutagenesis of *Trp53* and *Rb1* were noted only for group **F**. Group **B** had significantly less colonies than combined mutagenesis of *Trp53* and *Rb1* (Students t-test, $p<0.05$).

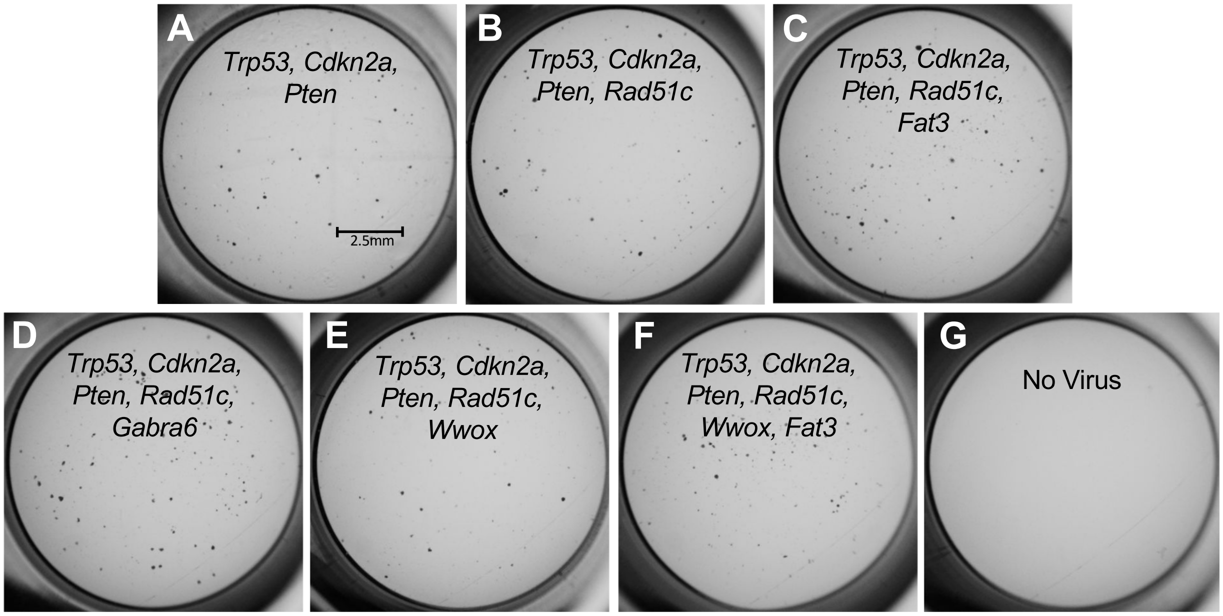


**Supplemental Figure 15. Combinatorial mutagenesis of Trp53, Cdkn2a, Pten, Rad51c and either Fat3 or Gabra6 causes significantly greater colony growth vs disruption of Trp53, CDkn2a, Pten, and Rad51c.** Cells were transduced with LentiCRISPRs targeting core genes *Trp53*, *Cdkn2a* and *Pten* **(A)** or *Trp53*, *Cdkn2a*, *Pten* and *Rad51c* **(B)** to establish a baseline degree of adhesion independent growth. OSE-SC were also transduced with LentiCRISPRs targeting *Trp53*, *Cdkn2a*, *Pten, Rad51c* and *Fat3* **(C)**, *Trp53*, *Cdkn2a*, *Pten, Rad51c* and *Gabra6* **(D)**, *Trp53*, *Cdkn2a*, *Pten, Rad51c* and *Wwox* **(E)**, *Trp53*, *Cdkn2a*, *Pten, Rad51c, Wwox* and *Fat3* **(F)** or no virus as a negative control **(G)**. Colonies were observed for all groups except the negative control. Significant increases in adhesion independent growth compared to cells transduced with *Trp53*, *Cdkn2a*, *Pten* and *Rad51c* were noted for groups **C** and **D** (Students t-test, $p<0.05$).

### Table S1: Primers for next generation sequencing

| Gene_exon_primer version | Sequence: illumina overhang_primer |
|---|---|
| BRCA1_4_1 | **TCGTCGGCAGCGTCAGATGTGTATAAGAGACAG**TTTTTCTGCTTATGCAGCATCT |
| BRCA1_4_1_rev | **GTCTCGTGGGCTCGGAGATGTGTATAAGAGACAG**AGTCAGAGGCCTTGTGCCTA |
| BRCA1_6_1 | **TCGTCGGCAGCGTCAGATGTGTATAAGAGACAG**CAGAGACCCTCCTGCTTCTG |
| BRCA1_6_1_rev | **GTCTCGTGGGCTCGGAGATGTGTATAAGAGACAG**AGAACACTTGTCCAGCCACT |
| BRCA1_7_1 | **TCGTCGGCAGCGTCAGATGTGTATAAGAGACAG**TTCATAGTTCTTTCCCTCTGTTCC |
| BRCA1_7_1_rev | **GTCTCGTGGGCTCGGAGATGTGTATAAGAGACAG**ACCAAAACTCCATGCAACAC |
| BRCA2_2_1 | **TCGTCGGCAGCGTCAGATGTGTATAAGAGACAG**GTAGCCACTTGGTTGGAAGC |
| BRCA2_2_1_rev | **GTCTCGTGGGCTCGGAGATGTGTATAAGAGACAG**GAGAAGTCCACCTGCCTCTG |
| BRCA2_3_1 | **TCGTCGGCAGCGTCAGATGTGTATAAGAGACAG**CCCCCATGACACAATAATGA |
| BRCA2_3_1_rev | **GTCTCGTGGGCTCGGAGATGTGTATAAGAGACAG**TTCTCTGAAAGGCGACTGGT |
| BRCA2_4_1 | **TCGTCGGCAGCGTCAGATGTGTATAAGAGACAG**GTGCACACCATCCCTTGAG |
| BRCA2_4_1_rev | **GTCTCGTGGGCTCGGAGATGTGTATAAGAGACAG**GCAAGATGACGGTTCATGACT |
| CSMD3_1_1 | **TCGTCGGCAGCGTCAGATGTGTATAAGAGACAG**CCGTTGGAGAGATGGTTGAG |
| CSMD3_1_1_rev | **GTCTCGTGGGCTCGGAGATGTGTATAAGAGACAG**AGCTTGCTAGCCTCTCTCAGG |
| FAT3_1_1 | **TCGTCGGCAGCGTCAGATGTGTATAAGAGACAG**TTGACCGACACACACCAGTT |
| FAT3_1_1_rev | **GTCTCGTGGGCTCGGAGATGTGTATAAGAGACAG**CTTTGTCTTTGGCCCTGACT |
| FAT3_2_1 | **TCGTCGGCAGCGTCAGATGTGTATAAGAGACAG**TGTTAATCTTCTCCTATTTGCCAAG |
| FAT3_2_1_rev | **GTCTCGTGGGCTCGGAGATGTGTATAAGAGACAG**AGTTCTGTGGGTTTCCACTTGT |
| FAT3_4_1 | **TCGTCGGCAGCGTCAGATGTGTATAAGAGACAG**TGAAATGGCCCAGTGAGTAA |
| FAT3_4_1_rev | **GTCTCGTGGGCTCGGAGATGTGTATAAGAGACAG**ACTCCCTGCTGTGAATTGCT |
| Gabra_2and2B | **TCGTCGGCAGCGTCAGATGTGTATAAGAGACAG**GGGATAGTGTGGCTGGTGTC |
| Gabra_2and2Brev | **GTCTCGTGGGCTCGGAGATGTGTATAAGAGACAG**TGGTCAGTGATTGAGGAAGG |
| GABRA_4_1 | **TCGTCGGCAGCGTCAGATGTGTATAAGAGACAG**TGCTGTCTTGATTAAATTTGGAA |
| GABRA_4_1_rev | **GTCTCGTGGGCTCGGAGATGTGTATAAGAGACAG**CAATCAGCGTTGATGGTGAG |
| RB1_1_2 | **TCGTCGGCAGCGTCAGATGTGTATAAGAGACAG**GGGGACGTTCCCATTATTTT |
| RB1_1_2_rev | **GTCTCGTGGGCTCGGAGATGTGTATAAGAGACAG**CTCCCTTCCCTTCCCTTCT |
| RB1_2_1 | **TCGTCGGCAGCGTCAGATGTGTATAAGAGACAG**AAAACTGTGCTGGTGTGTGC |
| RB1_2_1_rev | **GTCTCGTGGGCTCGGAGATGTGTATAAGAGACAG**CCACTGCCATCATCACCAT |
| RB1_4_1 | **TCGTCGGCAGCGTCAGATGTGTATAAGAGACAG**ATACTAGGGCCTGGGTTGCT |
| RB1_4_1_rev | **GTCTCGTGGGCTCGGAGATGTGTATAAGAGACAG**GCAAAATGGATAAGGCTAGGG |
| APC_4_1_4B_1 | **TCGTCGGCAGCGTCAGATGTGTATAAGAGACAG**GCAGGGCAAGTTTTAACTATTCT |
| APC_4_1_4Brev | **GTCTCGTGGGCTCGGAGATGTGTATAAGAGACAG**CCCACTCCCCTGTTACCTTT |
| APC_6_1 | **TCGTCGGCAGCGTCAGATGTGTATAAGAGACAG**TTTGGGTTTCTCAAGCATGG |
| APC_6_1_rev | **GTCTCGTGGGCTCGGAGATGTGTATAAGAGACAG**TAATGTCCAACAGCCACGAG |
| NF1_3_1 | **TCGTCGGCAGCGTCAGATGTGTATAAGAGACAG**AGCTTGGAAATTTATTTTTAGGG |
| NF1_3_1_rev | **GTCTCGTGGGCTCGGAGATGTGTATAAGAGACAG**TGGGATTTATAAAAGCTGAGAGAA |
| NF1_4_next | **TCGTCGGCAGCGTCAGATGTGTATAAGAGACAG**AGCTGTGTGGCTGTTCCTTC |
| NF1_4_next_rev | **GTCTCGTGGGCTCGGAGATGTGTATAAGAGACAG**CCTTCAAAAACCCAGATGTCA |
| LRP_1_1 | **TCGTCGGCAGCGTCAGATGTGTATAAGAGACAG**CTTTCGCTCACCTTCCACAT |

| | |
|---|---|
| LRP_1_1_rev | **GTCTCGTGGGCTCGGAGATGTGTATAAGAGACAG**GCACTCTGGCACCTAGTTCA |
| LRP_2_1 | **TCGTCGGCAGCGTCAGATGTGTATAAGAGACAG**GGAGTCAAGTCTGCCAAAGC |
| LRP_2_1_rev | **GTCTCGTGGGCTCGGAGATGTGTATAAGAGACAG**AGGCTTCCCATTTCTCTGGT |
| LRP_3_1 | **TCGTCGGCAGCGTCAGATGTGTATAAGAGACAG**TTTTTGGTGATCAGAACTGTGC |
| LRP_3_1_rev | **GTCTCGTGGGCTCGGAGATGTGTATAAGAGACAG**TTGAAGTTCATACTGAGAGACTGGA |
| PRIM_2_1 | **TCGTCGGCAGCGTCAGATGTGTATAAGAGACAG**TCAGCCTTTTTCCGTCATTT |
| PRIM_2_1_rev | **GTCTCGTGGGCTCGGAGATGTGTATAAGAGACAG**ATGGGCTGCTGAACAAAGAG |
| Prim_3_1 | **TCGTCGGCAGCGTCAGATGTGTATAAGAGACAG**CTGCGGAAACAAAGTTGTTAAT |
| Prim_3_1_rev | **GTCTCGTGGGCTCGGAGATGTGTATAAGAGACAG**GAGGGTGGGAAACTGCTGTA |
| Prim_4_1 | **TCGTCGGCAGCGTCAGATGTGTATAAGAGACAG**TGACTGATGGAAGGCAGTTG |
| Prim_4_1_rev | **GTCTCGTGGGCTCGGAGATGTGTATAAGAGACAG**ATATTCAGCCCATGGCACTC |
| CREBBP_2_1 | **TCGTCGGCAGCGTCAGATGTGTATAAGAGACAG**TTTTTCTTGTTTTACCTCCCTAAA |
| CREBBP_2_1_rev | **GTCTCGTGGGCTCGGAGATGTGTATAAGAGACAG**CCTAAGCTGGCCATGTTTGTA |
| CREBBP_3_1 | **TCGTCGGCAGCGTCAGATGTGTATAAGAGACAG**TGAGTGAAACTTGTCCTTTCACA |
| CREBBP_3_1_rev | **GTCTCGTGGGCTCGGAGATGTGTATAAGAGACAG**GCTTTTCACTGTGAGCACCA |
| Wwox_2_1 | **TCGTCGGCAGCGTCAGATGTGTATAAGAGACAG**CTGCCCTGCAAGATTCCTT |
| Wwox_2_1_rev | **GTCTCGTGGGCTCGGAGATGTGTATAAGAGACAG**GACATCCTTCTGTCCACCTCA |
| Wwox_3_1 | **TCGTCGGCAGCGTCAGATGTGTATAAGAGACAG**GAGTGCAACAGGGTTTTGGT |
| Wwox_3_1_rev | **GTCTCGTGGGCTCGGAGATGTGTATAAGAGACAG**AAGGGTGGAAAACTGCAGAC |
| Wwox_4_1 | **TCGTCGGCAGCGTCAGATGTGTATAAGAGACAG**TCCAGAGGCCAGAAGGTAGA |
| Wwox_4_1_rev | **GTCTCGTGGGCTCGGAGATGTGTATAAGAGACAG**GAGTCACCACCAACGTTTCA |
| Ankrd11_5_1 | **TCGTCGGCAGCGTCAGATGTGTATAAGAGACAG**ACTTCCCCTCATAGGCCTTC |
| Ankrd11_5_1_rev | **GTCTCGTGGGCTCGGAGATGTGTATAAGAGACAG**GAACCTCAAGACCCAACCAT |
| ANKRD11_6_1 | **TCGTCGGCAGCGTCAGATGTGTATAAGAGACAG**GGTCTTGAGGTTCCCATGAA |
| ANKRD11_6_1_rev | **GTCTCGTGGGCTCGGAGATGTGTATAAGAGACAG**CGCCCCCAATAAGAAAAAGA |
| ANKRD11_7_1 | **TCGTCGGCAGCGTCAGATGTGTATAAGAGACAG**ATTATGCAAAGCGCCACAAT |
| ANKRD11_7_1_rev | **GTCTCGTGGGCTCGGAGATGTGTATAAGAGACAG**CACAAAGCTCACTTCCCAGTC |
| MAP2K4_1_2 | **TCGTCGGCAGCGTCAGATGTGTATAAGAGACAG**CTCGGCTCTTCACTTCCAAC |
| MAP2K4_1_2_rev | **GTCTCGTGGGCTCGGAGATGTGTATAAGAGACAG**ACAGCGTTCACCGAAACC |
| MAP2K4_5_1 | **TCGTCGGCAGCGTCAGATGTGTATAAGAGACAG**CAAGCTAAAGGTTCAGCAGAGG |
| MAP2K4_5_1_rev | **GTCTCGTGGGCTCGGAGATGTGTATAAGAGACAG**TGAAGAATGTTCCTCAGATCCA |
| FancM_1_1 | **TCGTCGGCAGCGTCAGATGTGTATAAGAGACAG**AGAACGCTCTTCCAGACGTG |
| FancM_1_1_rev | **GTCTCGTGGGCTCGGAGATGTGTATAAGAGACAG**CCTGTCATTTCAGCCATGTG |
| FancM_2_1 | **TCGTCGGCAGCGTCAGATGTGTATAAGAGACAG**CCCGGCTTCACTGTTAATTT |
| FancM_2_1_rev | **GTCTCGTGGGCTCGGAGATGTGTATAAGAGACAG**CAGCACCCTCTGTTCTGACA |
| FancM_3_1 | **TCGTCGGCAGCGTCAGATGTGTATAAGAGACAG**ATGATTGGTGGTCGTTCCAT |
| FancM_3_1_rev | **GTCTCGTGGGCTCGGAGATGTGTATAAGAGACAG**ACCTTTAATCCCGGCACTTG |
| FancD2_2_1 | **TCGTCGGCAGCGTCAGATGTGTATAAGAGACAG**CATTTGGGTGGTTTGGACAG |
| FancD2_2_1_rev | **GTCTCGTGGGCTCGGAGATGTGTATAAGAGACAG**AAGCCCTAAGTGGCATTTGA |
| FancD2_5_1 | **TCGTCGGCAGCGTCAGATGTGTATAAGAGACAG**CTGTGGTGCTTCTGCTTCTG |
| FancD2_5_1_rev | **GTCTCGTGGGCTCGGAGATGTGTATAAGAGACAG**TTCCAGTTCTTTGCAATCCA |
| FancD2_6_1 | **TCGTCGGCAGCGTCAGATGTGTATAAGAGACAG**AACTCATAGCGGGCTGCTT |
| FancD2_6_1_rev | **GTCTCGTGGGCTCGGAGATGTGTATAAGAGACAG**TGGCAGATCAACCAGCCTAA |

| | |
|---|---|
| Rad51c_2_1 | **TCGTCGGCAGCGTCAGATGTGTATAAGAGACAG**CCATGCCAGGCTCTGAGTTA |
| Rad51c_2_1_rev | **GTCTCGTGGGCTCGGAGATGTGTATAAGAGACAG**TGTATGCTTCTTGGTTTCCTGA |
| Rad51c_3_1 | **TCGTCGGCAGCGTCAGATGTGTATAAGAGACAG**TTAGACATCTCTTTTGCCTTGG |
| Rad51c_3_1_rev | **GTCTCGTGGGCTCGGAGATGTGTATAAGAGACAG**TTTTGCAACAAAAGTATTGGAGA |
| Rad51c_5_1 | **TCGTCGGCAGCGTCAGATGTGTATAAGAGACAG**TCCAGCACCATCAAAAACTG |
| Rad51c_5_1_rev | **GTCTCGTGGGCTCGGAGATGTGTATAAGAGACAG**CCACAAGAACAACCACCAGA |
| Pten_1_1 | **TCGTCGGCAGCGTCAGATGTGTATAAGAGACAG**GAGAAGCAGGCCCAGTCTC |
| Pten_1_1_rev | **GTCTCGTGGGCTCGGAGATGTGTATAAGAGACAG**ATCTAGAAATGCGCCCAGAA |
| Pten_2_1 | **TCGTCGGCAGCGTCAGATGTGTATAAGAGACAG**AGTGAGTGGCTGACTGTCCA |
| Pten_2_1_rev | **GTCTCGTGGGCTCGGAGATGTGTATAAGAGACAG**CCAGTTCTCATCCAGTGACG |
| Pten_5_1 | **TCGTCGGCAGCGTCAGATGTGTATAAGAGACAG**TCTTACACTGGGATTATCTTTTTGC |
| Pten_5_1_rev | **GTCTCGTGGGCTCGGAGATGTGTATAAGAGACAG**CAGTTCTCAAAGCATCACACTG |
| CDK_2A_1 | **TCGTCGGCAGCGTCAGATGTGTATAAGAGACAG**CAACGTTCACGTAGCAGCTC |
| CDK_2A_1_rev | **GTCTCGTGGGCTCGGAGATGTGTATAAGAGACAG**CCTAGGCCTTGACCAGGAG |
| CDK_2B_1 | **TCGTCGGCAGCGTCAGATGTGTATAAGAGACAG**AGAATTCCAAGGCGGGACTA |
| CDK_2B_1_rev | **GTCTCGTGGGCTCGGAGATGTGTATAAGAGACAG**CAGCGGAACACAAAGAGCAC |
| CDK_2C_1 | **TCGTCGGCAGCGTCAGATGTGTATAAGAGACAG**GGCCGTGATCCCTCTACTTT |
| CDK_2C_1_rev | **GTCTCGTGGGCTCGGAGATGTGTATAAGAGACAG**GGGTTGCTTCTTCTTTTTCTGA |
| CSMD3_2_1 | **TCGTCGGCAGCGTCAGATGTGTATAAGAGACAG**TTTGTCTTGTGTTGCAGGATTT |
| CSMD3_2_1_rev | **GTCTCGTGGGCTCGGAGATGTGTATAAGAGACAG**TGAAAAACATATTCTTCGCATGA |
| CSMD3_4_1 | **TCGTCGGCAGCGTCAGATGTGTATAAGAGACAG**GGAGTTCCACCCAAAGGTGT |
| CSMD3_4_1_rev | **GTCTCGTGGGCTCGGAGATGTGTATAAGAGACAG**GTTATGCTTTATTTAACAGGTTTTCA |
| CREBBP_1_1 | **TCGTCGGCAGCGTCAGATGTGTATAAGAGACAG**GTGAAGATGGCCGAGAACTT |
| CREBBP_1_1_rev | **GTCTCGTGGGCTCGGAGATGTGTATAAGAGACAG**GCCTCTGGTCGCATTCTCT |
| MAP2K4_2_1 | **TCGTCGGCAGCGTCAGATGTGTATAAGAGACAG**AAGGCATGTTTCTCTCTTTTTCA |
| MAP2K4_2_1_rev | **GTCTCGTGGGCTCGGAGATGTGTATAAGAGACAG**ATCAGTCCCTTGCTTTGCAT |

**Legend:** Red/Green sequences are illumina overhangs

**Table S2: Primers for Surveyor assay**

| Gene_exon_primer version | Sequence: illumina overhang_primer |
|---|---|
| TRP53_4_1 | TTGTTTTCCAGACTTCCTCCA |
| TRP53_4_1_rev | GCATTGAAAGGTCACACGAA |
| TRP53_5_1 | TGGTGCTTGGACAATGTGTT |
| TRP53_5_1_rev | ACGCTGTGGCGAAAAGTCT |
| TRP53_6_1 | GCCATGGCCATCTACAAGAA |
| TRP53_6_1_rev | GGCAGCTTGCACCTCTAAG |
| RY_RB1_3_2 | GCTTACTCAAAGTTTCCAAAGGA |
| RY_RB1_3_2_rev | TGCACATGGTGCACTTACCT |
| RY_CSMD3_5_2 | TGAAATGATTAAATGTTTCTCACTGG |
| RY_CSMD3_5_2_rev | TCTCCCAGTTCTGACAATGC |
| RY_CSMD3_6_2 | TGAAGAATTCTACTTTGCAAGTTTGT |
| RY_CSMD3_6_2_rev | TTTGTAGCATGCTTTCTGCTG |
| RY_FancM_2_2 | ATAGAGCGAGCTAGCCGTGA |
| RY_FancM_2_2_rev | CAGCACCCTCTGTTCTGACA |
| RY_FancD2_8_2 | ATTTGGAAGAGCAGCCAGTG |
| RY_FancD2_8_2_rev | GGCTGTCCTGGAACTCACTC |
| RY_FancD2_9_2 | CCCCTGAACTCACACTCACA |
| RY_FancD2_9_2_rev | ATGAACAAGTTCCCCTGGTG |
| RY_PRIM2_6_2 | GCCCCATGTAGTGCTAGGAA |
| RY_PRIM2_6_2_rev | ATCCAGGAGCTTCAGCAATG |
| RY_BRCA2_5_2 | CATGAACCGTCATCTTGCAT |
| RY_BRCA2_5_2_rev | TACAGGCAGCTCACAACCAC |

**Table S3: Minilibrary sgRNA sequences**

| Minilib ID | Sequence | Exon |
|---|---|---|
| RY_TRP53_4_1 | AGTGAAGCCCTCCGAGTGTC | 4 |
| RY_TRP53_5_1 | GAAGTCACAGCACATGACGG | 5 |
| RY_TRP53_6_1 | AAATTTGTATCCCGAGTATC | 6 |
| RY_BRCA1_4_1 | ATTGTGAAGGCCCTTTCTTC | 4 |
| RY_BRCA1_6_1 | GGCGTCGATCATCCAGAGCG | 6 |
| RY_BRCA1_7_1 | GGTGTCCAGCTGTCTAACCT | 7 |
| RY_BRCA2_5F | TCTTACTCTGCGGTGCACAC | 5 |
| RY_BRCA2_3_1 | TAGGACCGATAAGCCTCAAT | 3 |
| RY_BRCA2_4_1 | GGACTAGCAACATCTACCAC | 4 |
| RY_CSMD3_1_1 | TTAGCGCATCTTCGCGTGCC | 1 |
| RY_CSMD3_5F | TTCATTAGGAAAACCCGGGC | 5 |
| RY_CSMD3_6F | GGTAGGGAGCACTAAATCCT | 6 |
| RY_FAT3_1_1 | GACCTTCACGCTGTAGCTAC | 1 |
| RY_FAT3_2_1 | AATAGAGAGGGACCACGCCC | 2 |
| RY_FAT3_4_1 | TTCTGGATGAAAACGACAAC | 4 |
| RY_GABRA_2_1 | TATGACAACCGTCTACGGCC | 2 |
| GABRA_2B_1 | AAGGCTATGACAACCGTCTA | 2 |
| RY_GABRA_4_1 | CCTGACACTTTTTTCCGGAA | 4 |
| RY_RB1_3F | TGTAGCTCAGTAAAAGTGAA | 3 |
| RY_RB1_2_1 | TTGGGAGAAAGTTTCATCCG | 2 |
| RY_RB1_4_1 | AGAAATCGATACCAGTACCA | 4 |
| RY_APC_4_1 | AGATCCTTCCCGACTTCCGT | 4 |
| RY_APC_4B_1 | GGATCTGTATCCAGCCGTTC | 4 |
| RY_APC_6_1 | GTGCACGCTTCTCCATGTCC | 6 |
| RY_NF1_3_1 | GTTGATCATATTGGATACAC | 3 |
| RY_NF1_4_1 | TCCGAAGTTCGGCTGCATGT | 4 |
| RY_NF1_4B_1 | ACCAACATGCAGCCGAACTT | 4 |
| RY_LRP_1_1 | GGGATTATTGCCTAACGCTG | 1 |
| RY_LRP_2_1 | TCGTGGCAAAGAAATTCGCC | 2 |
| RY_LRP_3_1 | GTCAAAACTGTGCAACGGAG | 3 |
| RY_PRIM_2_1 | CCGGAAGAAGCTGCGATTGG | 2 |
| RY_PRIM2_6F | CCATGATATCCTGTTCCCGA | 6 |
| RY_PRIM_4_1 | ATGAGTATGAGCCACGGCGA | 4 |
| CREBBP_1_1 | GTTGTCATTCGCGGAGAAGC | 1 |
| CREBBP_2_1 | CCATTGGGGATCAGCTCATC | 2 |
| CREBBP_3_1 | GGACAACCCTTTAGTCAAAC | 3 |
| RY_WWOX_2_1 | AACATCCGAAAACCGGCAAG | 2 |
| RY_WWOX_3_1 | TTTGCCGTATGGATGGGAAC | 3 |
| RY_WWOX_4_1 | CTTCGTCGGATTATCGTCCA | 4 |
| ANKRD11_5_1 | GTCCGGGCTGTTGTTCGGCA | 5 |

| | | |
|---|---|---|
| ANKRD11_6_1 | TGATGAGTTCCTTGATGCGC | **6** |
| ANKRD11_7_1 | TGGCGATGTCGTAATAGCCC | **7** |
| RY_MAP_1_1 | CCAGAAGCTGGAGGTCCGAT | **1** |
| RY_MAP_2_1 | CACCTGTCAAATCGACAGCA | **2** |
| RY_MAP_5_1 | TGTGTTAGATGACGTTATTC | **5** |
| RY_FancM_1_1 | GTCCAGCTGGTAGTCGCGCA | **1** |
| RY_FancM_2_1 | CACCCACGTAAAGTGTCTTG | **2** |
| RY_FancM_2F | TTACCATGACCTGCGGTGTC | **2** |
| RY_FancD2_8F | TAGTTGATTGATAATGAGTC | **8** |
| RY_FANCD_5_1 | GCTGTCTTGTGAGCGCCTGC | **5** |
| RY_FancD2_9F | ACTGCATCATCTGGGCCGTG | **9** |
| RY_RAD51_2_1 | TCTCGAGCAAGAGCATACCC | **2** |
| RY_RAD51_3_1 | GCCACGCCCCCAAAACATTC | **3** |
| RY_RAD51_5_1 | CATTTAGTAATCGAGTACGA | **5** |
| RY_PTEN_1_1 | GCTAACGATCTCTTTGATGA | **1** |
| RY_PTEN_2_1 | AAAGACTTGAAGGTGTATAC | **2** |
| RY_PTEN_5_1 | TGTGCATATTTATTGCATCG | **5** |
| RY_CDK_2A_1 | GTGCGATATTTGCGTTCCGC | **2** |
| RY_CDK_2B_1 | CGGTGCAGATTCGAACTGCG | **2** |
| RY_CDK_2C_1 | GGCTGGATGTGCGCGATGCC | **2** |
| Minilib 556 GFP1a | GAGCTGGACGGCGACGTAAA | **GFP** |
